# Enhancer RNAs stimulate Pol II pause release by harnessing multivalent interactions to NELF

**DOI:** 10.1101/2021.04.25.441328

**Authors:** Vladyslava Gorbovytska, Seung-Kyoon Kim, Filiz Kuybu, Michael Götze, Dahun Um, Keunsoo Kang, Andreas Pittroff, Lisa-Marie Schneider, Alexander Leitner, Tae-Kyung Kim, Claus-D. Kuhn

**Affiliations:** Gene Regulation by Non-coding RNA, Elite Network of Bavaria and University of Bayreuth, Universitätsstrasse 30, 95447 Bayreuth, Germany; Department of Life Sciences, Pohang University of Science and Technology (POSTECH), Pohang, Gyeongbuk, 37673, Korea; Department of Biology, Institute of Molecular Systems Biology, ETH Zurich, 8093 Zurich, Switzerland; Department of Microbiology, Dankook University, Cheonan, 31116, Korea

**Author notes:** These authors contributed equally to this work (Vladyslava Gorbovytska, Seung-Kyoon Kim). Correspondence should be addressed to: T.-K. Kim, C.-D. Kuhn.

## Abstract

Enhancer RNAs (eRNAs) are long non-coding RNAs that originate from enhancers. Although eRNA transcription is a canonical feature of activated enhancers, the molecular features required for eRNA function and the mechanism of how eRNAs impinge on target gene transcription have not been established. Thus, using eRNA-dependent RNA polymerase II (Pol II) pause release as a model, we examined the requirement of sequence, structure and length of eRNAs for their ability to stimulate Pol II pause release by detaching NELF from paused Pol II. We found eRNA not to exert their function through common structural or sequence motifs. Instead, efficient NELF release requires a single eRNA molecule that must contain unpaired guanosines to make multiple, allosteric contacts with several NELF subunits. By revealing the molecular determinants for eRNA function, our study mechanistically links eRNAs to Pol II pause release and provides new insight into the regulation of metazoan transcription.

## INTRODUCTION

Enhancers are DNA elements that direct metazoan tissue- and stimulus-specific gene expression by serving as transcription factor binding platforms. On primary sequence level enhancers can be distant from their target genes; however, to achieve regulation, enhancers and promoters are brought into proximity by chromatin looping^1^. The regulatory information encoded in the composition of the enhancer-bound transcription factors is transmitted to the core transcription machinery via Mediator^2^. In addition to orchestrating transcription factor binding, enhancers themselves are transcribed bidirectionally by RNA polymerase II (Pol II), resulting in the production of large numbers of long non-coding RNAs (lncRNAs) that are termed enhancer RNAs (eRNAs)^3,4^. In general, eRNA expression has been shown to correlate with target gene activation, but whether eRNA molecules themselves are functional or whether it is the act of transcription that boosts enhancer activity is currently under debate^5–8^.

Although considerable research has focused on histone modification and transcription factor binding profiles at enhancers, recent studies show that eRNA levels and the extent of their bidirectional transcription are most likely a better predictor of enhancer activity^9,10^. Despite their inherent instability^11^, eRNAs were detected in different tissues and following numerous stimuli^3–5,12,13^, and they were shown to correlate with the activation of diverse sets of target genes. This data strongly supports a direct, functional role for eRNAs, but their mechanistic role remains largely unexplained. This is in part due to the fact that eRNAs, similar to other lncRNAs, possess diverse mechanisms through which they exert their function. For example, in cell culture experiments eRNAs were shown to increase transcription rates by stimulating the histone acetyltransferase activity of CBP (CREB-binding protein)^14^. Furthermore, eRNAs cause the increased retention of transcription factors, such as YY1 and BRD4, at gene regulatory elements, thereby modulating their biological impact^15,16^. In earlier work, we characterized the impact of eRNAs on the stimulus-induced expression of immediate early genes (IEGs) in mouse primary neurons^17^. Our results suggested that neuronal eRNAs trigger the release of NELF from paused Pol II at IEGs, whereby they presumably facilitated the transition of Pol II into productive elongation. To achieve NELF release, we speculated that eRNAs might compete with nascent mRNAs for binding to negative elongation factor (NELF). However, the molecular mechanism of how eRNAs affect paused Pol II and the structural and sequence characteristics that enable eRNA-driven NELF release remained unknown.

Here we answer these unsolved questions and reveal an unexpected allosteric mechanism through which eRNAs stimulate PoI II pause release. First, we characterize both the sequences and the structures of dozens of neuronal eRNAs by a combination of Exo-seq (5’-end RNA-seq) and SHAPE-MaP (selective 2’-hydroxyl acylation analyzed by primer extension and mutational profiling). Second, we then use electrophoretic mobility shift assays (EMSAs) to show that eRNAs efficiently trigger NELF release from the paused elongation complex (PEC) *in vitro*, but only if eRNAs are sufficiently long and contain several unpaired guanosines. Third, we apply protein-RNA crosslinking coupled to mass spectrometry on the eRNA-bound PEC and uncover that the length requirement for eRNA functionality stems from multiple RNA binding sites on the NELF complex and on the PEC. In order to detach NELF from Pol II, eRNAs must bind several RNA binding sites simultaneously. These *in vitro* findings are supported by NELF-E-directed eCLIP-seq (enhanced UV crosslinking and immunoprecipitation sequencing) experiments in mouse primary neurons, which demonstrate that NELF directly contacts eRNAs primarily in their 5’-terminal 200 nucleotides (nt). By utilizing a reconstituted pause release assay, we further demonstrate that eRNA-driven NELF release results in transcription activation through the more efficient release of Pol II from the paused state. Complementing these *in vitro* findings once more *in vivo*, we find that NELF binding levels correlate with rapid and efficient transcriptional elongation in response to neuronal stimulation, suggesting that eRNA-dependent release of NELF from paused Pol II could play a role in activity-induced transcription. To our knowledge this study represents the first report that mechanistically links eRNAs to the core Pol II transcription machinery. Furthermore, by revealing the detailed molecular determinants that enable eRNAs to stimulate Pol II pause release, our study establishes the functional capacity of eRNAs in the regulation of metazoan transcription.

## RESULTS

### Exo-seq allows for the assignment of eRNA transcription start sites with single-nucleotide precision

To begin to decipher the molecular mode of action of eRNAs in mouse primary neurons, it was imperative to first determine their exact sequences. We thus performed global run-on sequencing (GRO-seq)^18^ before and after neuronal stimulation by KCl treatment and profiled all resulting nascent RNAs (see Methods). We defined transcription units *de novo* from GRO-seq reads and assigned them either to annotated genes or to eRNAs based on the overlap between intergenically-located transcript units and regions enriched for histone H3 lysine 27 acetylation (H3K27ac), a histone mark previously found to label active enhancers^19^. A total of 9,028 annotated genes were overlapped with the transcript units defined *de novo* from GRO-seq reads. In addition, we identified a total of 1,226 intergenic eRNA transcription units, of which 252 were activity-induced, as defined by a >1.5-fold increase of eRNA GRO-seq signal at any time point after KCl stimulation (Supplementary Table 1). As our GRO-seq data did not allow us to unambiguously determine the 5’-ends of eRNAs, we performed the Exo-seq protocol in addition, a technique dedicated to the assignment of RNA 5’-ends^20^. Indeed, after applying the program TSScall^10^ to our Exo-seq data, we could determine the 5’-ends of eRNAs with single-nucleotide precision. We then intersected the list of eRNA TSSs (eTSS = enhancer transcription start site) with our GRO-seq-based list of 1,226 intergenic eRNA transcription units to find 304 of them (281 for replicate 2) to exhibit well-defined 5’-ends (>20 reads per eTSS). 86 of these eRNAs (79 for replicate 2) stem from activity-induced enhancers. Overall, the 5’-ends of both eRNAs and mRNAs - as defined by Exo-seq - were well-correlated with nascent transcripts detected by GRO-seq, as shown for two enhancer loci, *Nr4a1* and *Arc* (Fig. 1a,b). In corroborating our analysis, we detected eRNA TSSs of IEGs such as *Arc*, *Nr4a1*, *Junb*, *c-Fos* (enhancers e1, e2 and e5) and *Fosb*, for some of which eRNA expression had been reported before^3^. After closer inspection of the Exo-seq reads at the selected eTSSs, we excluded all sites with pervasive or convergent transcription and compiled a final list of 33 high-quality eRNA candidates, including all abovementioned IEG eRNAs. Seven of the 33 eRNA candidates, amongst them *Junb and Nr4a1*, showed distinct, likely alternative eTSSs. For these eRNAs we therefore included two separate eRNAs (termed (a) and (b)). In total, this resulted in a test set of 39 eRNAs (Fig. 1b and Supplementary Table 2).

**Fig. 1.**
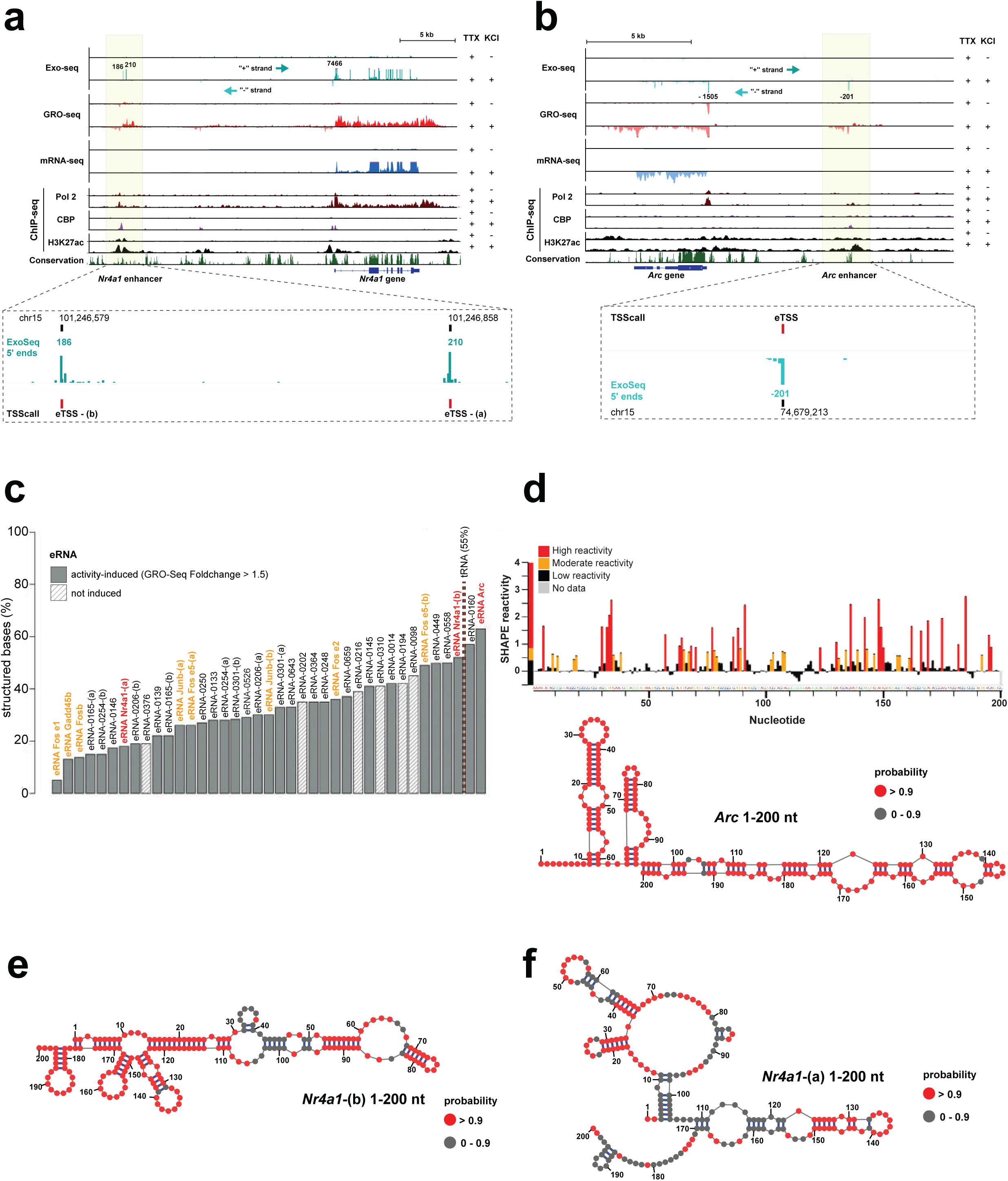
Secondary structure mapping of selected mouse neuronal enhancer RNAs. The mouse *Nr4a1* **(a)** and *Arc* **(b)** gene and enhancer loci in genome browser view. Exo-, GRO-, and mRNA-seq signal from KCl-stimulated mouse cortical neurons is plotted alongside Pol II (8WG16 antibody), CBP and H3K27ac ChIP-Seq data from publicly available sources3,19. A zoom-in view of the Exo-seq data attests to the single nucleotide resolution in determining eRNA TSSs (eTSS). The enhancer locus of *Nr4a1* revealed two distinct TSS sites (eTSS-(a) and eTSS-(b)). **c,** Percentage of structured bases (base pairing probability p >=0.9) for all 39 eRNAs in our data set. eRNAs possess both low and high percentages of structured bases (shown from left to right side of the plot). **d,** SHAPE reactivity profile of *Arc* eRNA (1-200), as computed by using *Shapemapper2*. Using the experimentally determined SHAPE reactivities as restraints, the corresponding secondary structure (shown below) was generated with *RNAStructure*27. High (>=0.85, red) and moderate (>= 0.4, orange) SHAPE reactivities mark flexible, unpaired nucleotides. Low SHAPE reactivities (< 0.4, black) denote less-flexible nucleotides that are predominantly located in double-stranded regions. Nucleotides are shown as circles, and they are colored in red if their base pairing probability exceeds 0.9. **(e** and **f)** Secondary structures of variant (b) and variant (a) of *Nr4a1* eRNAs (1-200). Determined and plotted as in (d).

### RNA structure probing uncovers that eRNAs do not share common structural motifs

As we previously showed that target gene induction depends on eRNAs^17^, yet we were unable to identify sequence motifs that would explain eRNA function, we speculated that eRNA might possess specific structural features that are important for their function. We thus set out to determine the secondary structures of all 39 eRNA candidates, for which we had identified the precise eTSS (Fig. 1a,b and Supplementary Table 2). Although eRNAs are reported to have a median length of ∼ 1kb^11^, we decided to focus our secondary structure analysis on the 5’-terminal 200 nt of all eRNAs for the following reasons: 1. The few already experimentally determined lncRNA structures hint at minimal lncRNA domains of approximately 200 nucleotides^21^, and 2. eRNAs are capped and thus possess defined 5’-ends, whereas their 3’-ends are not polyadenylated and therefore prone to exosome-mediated decay^22^. The functionally relevant eRNA parts are therefore likely confined to their 5’-ends. To map all 39 eRNA structures simultaneously, we turned to chemical probing of RNA secondary structure read out by next-generation sequencing. Specifically, we performed the SHAPE-MaP protocol with *in vitro* transcribed^23^, monodisperse eRNA (1-200) fragments (Supplementary Fig. 1a) by use of 1M7 (1-methyl-7-nitroisatoic anhydride) as a modifying agent^24,25^. Following sequencing of our SHAPE-MaP libraries, we calculated mutation rates and SHAPE reactivities using ShapeMapper2^26^ and computed SHAPE-restraint RNA secondary structures using RNAstructure^27^ (Supplementary Fig. 1b,c and Supplementary Table 3). Next, we quantified the “structuredness” of all eRNAs by focusing on the number of base pairs that form with a probability of >0.9 (Fig. 1c). To our surprise, this analysis revealed that our set of eRNAs folds into a wide range of structures. On average, 32% of all nucleotides exhibit a base pairing probability > 0.9, however, overall, base pairing ranges from 5% of all nucleotides for *Fos* e1 eRNA (1-200) to 63% for *Arc* eRNA (1-200) (Fig. 1c). As tRNAs display 55% of paired nucleotides, we consider *Arc* eRNA (1-200) to be highly structured (Fig. 1d). We did not find a high degree of structuredness to be a characteristic of eRNAs, a finding that can be rationalized by analyzing the two alternative *Nr4a1* eRNA variants (Fig. 1e,f). Whereas variant (b) shows a portion of base-paired nucleotides that is comparable to *Arc* eRNA (52%), variant (a) only encompasses 18% of paired nucleotides. In summary, our experimental data on the structures of the 5’-terminal 200 nt of 39 neuronal eRNAs strongly argue against a general lack of eRNA structuredness, as was reported before using RNA fold predictions^11^. In contrast, we find eRNAs to populate a wide range of structural space without any common structural motifs.

### NELF detachment from the paused elongation complex is markedly dependent on eRNA length

The diverse nature of their structures did not offer a path forward in deciphering the mechanistic impact of eRNAs on paused Pol II (Fig. 1 and Supplementary Table 3). To remedy this, we reconstituted the mammalian paused elongation complex (PEC) *in vitro*^28,29^. To then study the effect of eRNAs on the PEC, we used electrophoretic mobility shift assays (EMSAs), as this technique offers both high sensitivity for detecting protein-RNA interactions and allows for the study of large macromolecular complexes such as the PEC (which has a size of 0.9 MDa). The PEC was stepwise assembled on a synthetic transcription bubble (Fig. 2a) that contained a 5’-³²P-labelled “nascent” RNA with a length of 25 nt^28,29^. Our EMSA setup confirmed that nascent RNA needs to be >22 nt in length to allow for PEC assembly (Supplementary Fig. 2a)^30^. Following PEC assembly, we added the 5’-terminal 200 nt fragments of *Arc*, *Nr4a1*-(a) or *Nr4a1*-(b) eRNAs, respectively, and analyzed the mobility of formed complexes (Fig. 2b-d, and Supplementary Fig. 2b). Intriguingly, at eRNA concentrations only about equimolar to PEC concentrations, we already observed NELF dissociation from the PEC (top gels in Fig. 2b-d). In contrast to NELF, DSIF (DRB sensitivity-inducing factor) remained untouched and stably bound to Pol II after eRNA addition - even in presence of a 10-fold excess of eRNA (top gels in Fig. 2b-d). Our results are in agreement with established knowledge on pause release and they assert the validity of our assay setup for studying pausing *in vitro*^31–33^. We also confirmed the presence of NELF and DSIF by performing supershift assays (Supplementary Fig. 2c). Next, to uncover the part or parts of each eRNA that are responsible for NELF dissociation, we shortened the eRNAs to their 1-100 and 1-50 variants (middle and lower gels in Fig. 2b-d). When using these shorter variants, we were very surprised to find that, whereas *Arc* eRNA and *Nr4a1*-(b) (1-100) fragments retained some of their NELF-dissociation potential, both 1-50 fragments were unable to induce NELF dissociation from the PEC, even in 18-fold molar excess (middle and lower gels in Fig. 2b-d). To assess the dissociation potential of different eRNA fragments towards NELF, we quantified the appearance of the Pol II-DSIF complex following NELF dissociation (Supplementary Fig. 2e). Our analysis confirmed that eRNA (1-100) fragments exhibit a > 10x higher apparent K_d_ compared to their 1-200 siblings and, thus, possess a clearly reduced ability to displace NELF. In contrast to the *Nr4a1*-(b) (1-100) fragment, the *Nr4a1*-(a) (1-100) fragment was considerably more effective in dissociating NELF (K_d_=2.02 µM vs. 0.14 µM) (Fig. 2d,e). Further, the *Nr4a1*-(a) 1-50 fragment also displayed a mild NELF dissociation effect with a K_d_ of 9.70 µM. Interestingly, while *Arc* and *Nr4a1*-(b) are highly structured with prominent double-stranded regions (Fig. 1d,e), *Nr4a1*-(a) exhibits a more flexible structure with long single-stranded regions (Fig. 1f). This hints at the possibility that RNA flexibility might facilitate the observed dissociative effect of eRNAs on the PEC. Taken together, we found eRNA-induced NELF dissociation to be critically dependent on eRNA length. However, due to their divergent structures and sequences, these results did not reveal the mechanism of eRNA-induced NELF dissociation.

**Fig. 2.**
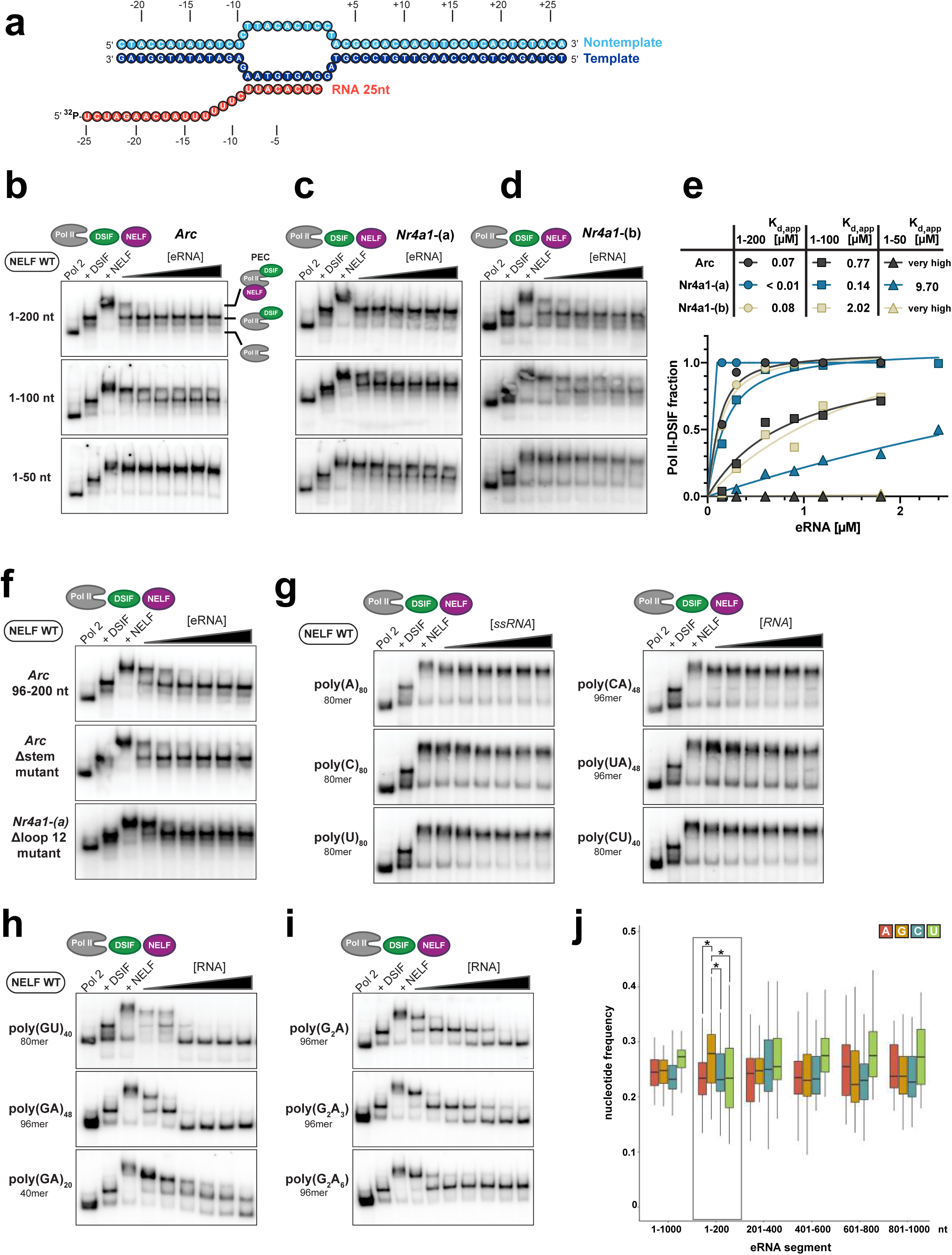
eRNAs trigger NELF release from the paused elongation complex in a length- and guanosine-dependent manner *in vitro*. **a,** Nucleic acid scaffold used for radioactive electrophoretic mobility shift assays (EMSAs). Template DNA is shown in dark blue and has a length of 49 nt, long fully-complementary non-template DNA is shown in light blue and is 49 nt long, nascent RNA has a length of 25 nt and is shown in red. **b,** EMSAs demonstrate the ability of *Arc* eRNA variants to release NELF from the PEC (paused elongation complex). EMSAs using *Arc* eRNAs with three different lengths are shown: The top gel show experiments with Arc eRNA (1-200), the middle gel examines the effect of Arc eRNA (1-100), the bottom gel shows the effect for Arc eRNA (1-50) on the PEC. **c,** EMSAs with the *Nr4a1*-(a) variant. Experimental setup and display as described for (b). **d,** EMSAs with the *Nr4a1*-(b) variant. Experimental setup and display as in (B). For all EMSAs the PEC was assembled on the nucleic acid scaffold shown in (a) using 0.8 pmol 32P-labeled RNA. The Pol II-DSIF complex was then formed by using 1.2 pmol Pol II and 2.4 pmol DSIF. Subsequently, 1.2 pmol NELF were added to form the PEC (final concentration 0.1 µM). The addition of increasing amounts of eRNAs (final concentrations: 0.15, 0.3, 0.6, 0.9, 1.2, 1.8, 2.4 µM) triggered detachment of NELF from the PEC. **e,** NELF release from the PEC is quantified by plotting the formation of the Pol II-DSIF complex against the eRNA concentration (mean of two experimental replicates). The resulting pseudo binding curve was fitted using a single-site binding model, apparent Kd values are indicated. **f,** EMSAs performed with the highly-structured *Arc* eRNA (96-200) fragment (length = 104 nt), a mostly single-stranded *Arc* eRNA Δstem mutant (length = 100 nt) and a *Nr4a1*-(a) Δloop 12 mutant (length = 102 nt). The *Nr4a1*-(a) Δloop 12 mutant lacks prominent single-stranded regions present in *Nr4a1*-(a) eRNA (see Supplementary Fig. 2d). EMSAs were carried out as described for (B-D). **g,** EMSAs performed with single-stranded, low complexity RNAs, such as poly (A), poly (C), poly (U), poly (CA), poly (UA), and poly (CU) RNAs. **h,** EMSAs performed with 80 nt long poly (GU) RNA and different lengths of poly (GA) RNA (40 and 96 nt). **i,** EMSAs performed with 96 nt long poly (G_2_A), (G_2_A_3_) and (G_2_A_6_) RNA. **j,** Nucleotide frequency plot for all 39 experimentally verified eRNAs (Fig. 1c and Supplementary Table 2). eRNA sequences were extended to 1 kb and divided into bins of 200 nt. Only in the 5’-terminal 200 nt are guanosines significantly overrepresented (as determined by a pairwise t-test with p-values (A/G) = 0.025; (C/G) =0.029; (U/G) =0.029).

### Unpaired guanosines are critical for the dissociative effect of eRNAs on the paused elongation complex

To further decode the functional role of eRNA structure, we analyzed the almost entirely double-stranded *Arc* eRNA (96-200) fragment and an *Arc* mutant that lacked almost all secondary structure (Arc Δstem 1-100) (Supplementary Fig. 2d). Despite their diametrically opposed structures, both eRNAs were able to detach NELF from Pol II, clearly demonstrating that eRNA function does not depend on structure alone (Fig. 2f). Thus, to simplify the sequence and structure space of the tested RNAs, we turned to synthetic, low complexity RNAs and measured their effect on NELF dissociation. Surprisingly, none of the tested single-stranded RNAs that lack guanosines showed the ability to dissociate NELF from the PEC (Fig. 2g). In contrast, despite their short length G-containing poly (GU)_40_ RNA and poly (GA)_48_ RNA were able to displace NELF at low RNA concentrations and even led to DSIF dissociation at higher RNA concentrations (Fig. 2h). As observed before (Fig. 2b-e), we found this effect to be dependent on RNA length. Next, we asked which “density” of guanosines is sufficient to trigger NELF release, and thus we tested synthetic 96-mer poly (G_2_A) RNA, poly (G_2_A_3_) RNA and poly (G_2_A_6_) RNA. As for poly (GU)_40_ RNA and poly (GA)_48_ RNA, we found all of them to be able to fully dissociate NELF. However, at higher RNA concentrations poly (G_2_A_6_) RNA, bearing the widest G spacing, did not cause DSIF dissociation (Fig. 2i). In summary, RNA-driven NELF dissociation from the PEC seems critically dependent on unpaired guanosines. In building on this, we were able to rationalize our prior data on eRNA fragments (Fig. 2b-e). There, we only found the short fragment of *Nr4a1-*(a) eRNA to induce the partial release of NELF (Fig. 2c,e). This effect can now be explained by focusing on the number of unpaired guanosines that do not form stable base pairs with cytidines. Despite the fact that, overall, *Arc* eRNA (1-50) and *Nr4a1*-(b) eRNA (1-50) contain more guanosines, these are mostly paired with cytidines, which is not the case for *Nr4a1*-(a) (1-50) (Supplementary Fig. 2e). To experimentally substantiate the functional role of unpaired guanosines, we generated G-less *Nr4a1*-(a) eRNA (1-50) and (1-100) variants, in which we replaced all guanosines with adenosine (G-to-A mutant) or cytidine (G-to-C mutant). Indeed, the G-less mutants showed a strongly reduced ability to trigger NELF release, as reflected by 10x higher apparent K_d_ values (Supplementary Fig. 2f). Interestingly, the restoration of two guanosines in the middle (2G middle) or three guanosines at the 3’- or at the 5’-end of the G-less mutant (3G at 5’-end or 3G at 3’-end) enhanced eRNA-driven NELF detachment 2.5 to 5-fold, respectively (K_d_ = 0.38 µM for 3G at the 5’-end and K_d_ = 0.32 µM for 3G at the 3’-end) (Supplementary Fig. 2f). However, the additional restoration of three guanosines at the RNA’s 5’- or 3’-end (6 guanosines in total) did not facilitate NELF dissociation any further. As the apparent K_d_ for NELF release by WT *Nr4a1*-(a) (1-100) eRNA is lower the any of the mutants with restored guanosines (0.14 µM, Fig. 2e), our data argue that potent eRNAs must comprise several widely spaced unpaired guanosines and not only one cluster. Last, we sought to support our *in vitro* data by a more global analysis on the prevalence of guanosine in eRNAs. Thus, we calculated the nucleotide distribution across our set of 39 neuronal eRNAs (Supplementary Table 2). Indeed, we found guanosine to be overrepresented in their first 200 nt in a statistically significant manner (Fig. 2j), but not amongst the larger collection of eRNAs (Supplementary Table 1). This suggests that only a subset of eRNAs (*e.g.*, activity-induced eRNAs) possess elevated levels of guanosine within their first 200 nt. Intriguingly, we only find guanosine to be overrepresented, which sets our finding apart from 5’-UTRs of coding genes. These are highly structured and hence show a higher content of both guanosine and cytidine^34^.

### eRNAs bind to a positive patch on NELF-C and to the NELF-E RRM domain

For eRNAs to displace NELF from the PEC they must very likely directly contact NELF, a complex of four proteins termed NELF-A, -B, -C and -E^35^. The RRM (RNA recognition motif) domain of NELF-E was shown to bind both single-stranded, as well as structured RNAs, such as the HIV TAR element, *in vitro*^36–39^. Previously, we had shown that the RRM domain is essential for eRNA function *in vivo*^17^. Moreover, RNA binding studies with recombinant NELF had identified two additional RNA binding sites on NELF-A/C and on NELF-B^40^. Whereas the NELF-E RRM domain might be involved in contacting nascent RNA during pausing^38^, the biological relevance of the other two RNA binding interfaces has not been established. To decipher how eRNAs are able to dissociate NELF from paused Pol II, we therefore purified a NELF variant that lacked the RRM domain of NELF-E (NELFΔRRM) and examined its dissociation from the PEC after eRNA addition. All mutant PEC complexes could be assembled as efficiently as wild-type complexes (Supplementary Fig. 3a), but we observed significantly diminished NELF dissociation induced by *Arc* eRNA (1-200 and 1-100) fragments (K_d_s of 0.33 µM and 7.8 µM), suggesting that the RRM domain of NELF-E is directly involved in eRNA-driven NELF release (Fig. 3a). We detected an analogous effect for *Nr4a1*-(b) eRNA fragments (Supplementary Fig. 3b). However, the absence of the NELF-E RRM domain did not fully abolish NELF release, a finding which was particularly obvious for the eRNA 1-200 nt variants and for *Nr4a1*-(a) (1-100) eRNA (Fig. 3a,f and Supplementary Fig. 3b). This demonstrates that eRNA-driven dissociation of NELF from the PEC does not solely depend on the RRM domain. It is also in line with the observed dependency of NELF dissociation on eRNA length. To uncover the missing eRNA binding site on NELF, we next turned to a NELF mutant, in which one of the two additional RNA binding sites, a positively charged surface patch on NELF-A/C had been mutated (NELF-C patch mutant)^40^. Using this mutant, we found dramatically diminished eRNA-driven NELF dissociation from the PEC (Fig. 3b,f and Supplementary Fig. 3c). More specifically, *Arc* and both *Nr4a1* eRNA (1-100) fragments hardly induced NELF detachment, whereas the 1-200 nt fragments did, but to a much lesser extent as compared to the NELFΔRRM variant. Last, we combined both mutants to perform EMSAs with a NELF variant that lacked both the NELF-E RRM domain and whose NELF-C patch was mutated (Fig. 3c-e). Intriguingly, this double mutant has entirely lost its ability to dissociate from the PEC, even if eRNA (1-200) fragments were added in large molar excess. Thus, we conclude that both the NELF-E RRM domain and the positively charged surface patches on the NELF AC-lobe are essential to enable eRNA-induced NELF dissociation from the PEC (Fig. 3).

**Fig. 3.**
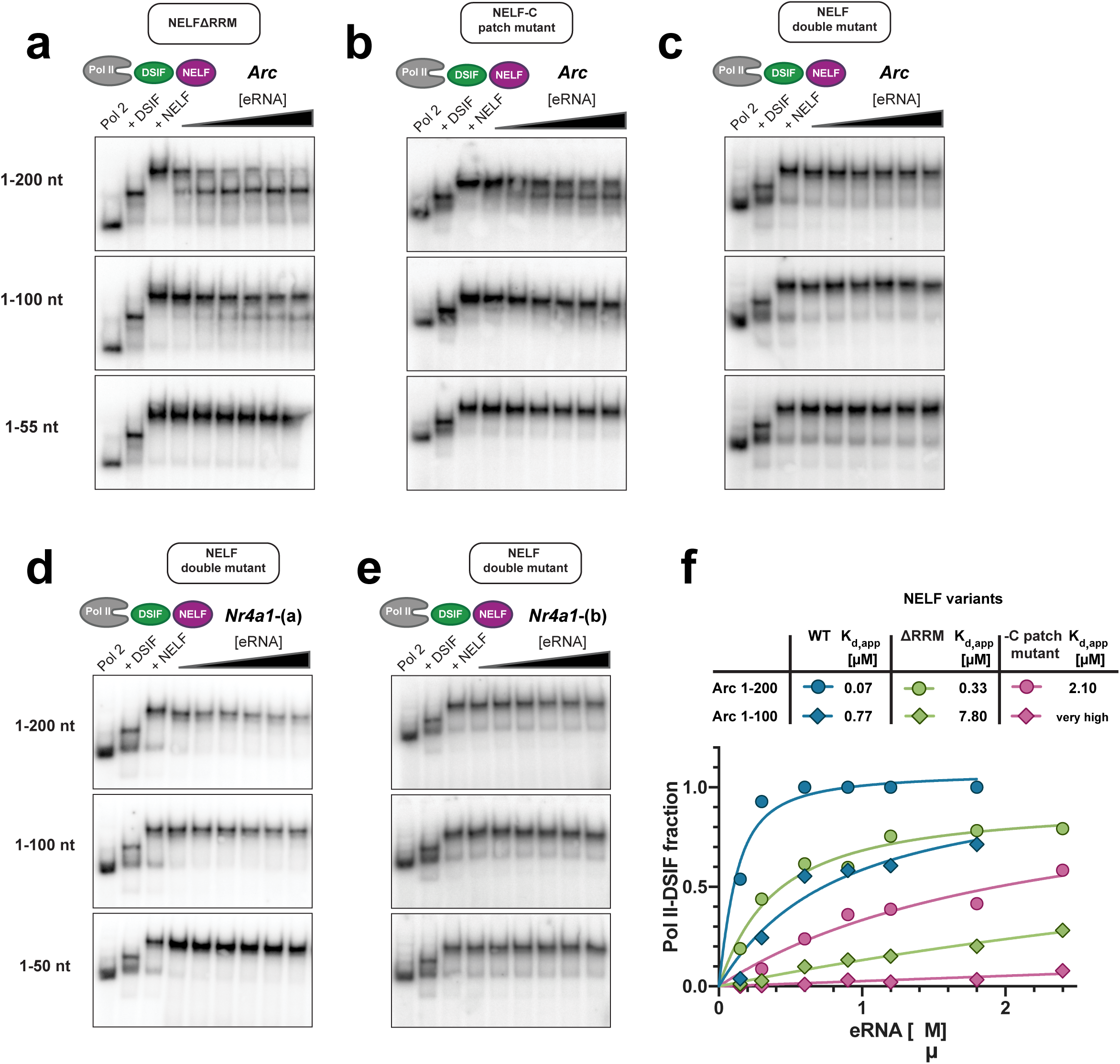
Positive patches on NELF-AC and the NELF-E RRM domain are both involved in eRNA-driven NELF dissociation. EMSAs carried out with PEC variants that comprise **(a)** the NELF-ABC-EΔRRM complex (NELF ΔRRM), **(b)** a mutant NELF-ABCE complex, in which positively charged surface patches on NELF-C are removed (NELF-C patch mutant)40, or **(c)** a NELF double mutant (NELF patch mutant + NELF-E ΔRRM). All assays were performed as in Fig. 2b-d, using *Arc* eRNAs (1-50), (1-100) or (1-200). In **(d)** and **(e)** EMSAs of the NELF double mutant using *Nr4a1*-(a) and *Nr4a1*-(b) eRNA fragments are shown, respectively. **f,** Quantification of NELF release using data from Fig. 3a-c. Data for wild-type NELF (Fig. 2b) are shown to aid comparison. Binding curves were plotted and fitted as in Fig. 2e.

### Protein-RNA crosslinking coupled to mass spectrometry uncovers an extended interaction network of eRNAs and the Pol II paused elongation complex

To corroborate our hypothesis that eRNAs simultaneously bind to NELF-C and the NELF-E RRM domain to trigger NELF release from the PEC, we turned to UV-induced protein-RNA crosslinking followed by mass spectrometry. We assembled the PEC as described^28^ and then further added *Arc* eRNA (1-200) or *Nr4a1*-(b) eRNA (1-100). We then separated the eRNA-bound PEC from non-incorporated components by size-exclusion chromatography (Supplementary Fig. 4a). The presence of eRNA in our PEC preparation was confirmed by both urea PAGE analysis and by an increased A260/A280 absorbance ratio as compared to the regular PEC assembly (Supplementary Fig. 4a,b). Under our experimental conditions only a fraction of the PEC is bound to an eRNA, as reflected by the relatively small increase of A260 absorbance after addition of *Arc* eRNA (1-200), which has a size of 66 kDa (33 kDa for *Nr4a1*-(b) (1-100)). This low binding efficiency is likely due to our ability to isolate transient eRNA-PEC complexes that, at higher eRNA concentrations, would trigger the complete dissociation of the PEC, as demonstrated in Fig. 2b-d. After UV crosslinking, limited RNase digestion and subsequent tandem mass spectrometry analysis, we retrieved 83 unique protein-RNA crosslinks for the *Arc* eRNA (1-200)-bound complex and 237 crosslinks for the *Nr4a1*-(b) eRNA (1-100) variant (Supplementary Table 4). Our data did not allow us to unambiguously detect >2 nt attached to any amino acid, which precluded the unambiguous assignment of RNA sequence to protein surface. However, we obtained highly scoring protein-RNA crosslinks between uridines and residues of the RPB1 dock domain of Pol II, the RPB2 wall, switch 3 loop region and the SPT5 KOWx-4 and KOW4-5 linker region (Supplementary Fig. 4c). As these residues were shown to contact nascent RNA within the Pol II-DSIF and the PEC structures, we can unambiguously attribute them to a stretch of uridines in nascent RNA (position −9 through −15), thereby confirming the integrity of our PEC preparation (Fig. 4a)^28–30,41^. Next, and most interestingly, our data confirm RNA binding to NELF (Fig. 4b). In focusing on the NELF-AC lobe, we found numerous polar residues on NELF-A (S71, T173, Q180, T185) and on NELF-C (S301, K302, K311, S377, L411, L414) to be crosslinked to RNA. These residues are located in vicinity of the positively charged patches 1, 2 and 4 that are part of the NELF-AC dimer (detailed view in Fig. 4c). These patches were demonstrated to impact RNA binding to NELF^40^ and we found them responsible for eRNA-dependent NELF dissociation from the PEC (Fig. 3b-e). Moreover, we observed multiple RNA crosslinks to the unstructured N-terminal domain of NELF-E (Q35, T40, S42, Q43, K47, A59, T60, I74, E93, L96, K97, D98, E134, V137), along with crosslinks to the NELF-E tentacle just before its RRM domain (S249 – P253) and within the RRM domain (D288, L289, D292, K304) (Supplementary Fig. 4d). Intriguingly, many of the RNA-protein crosslinks lie in proximity to protein-protein crosslinks that had been reported between NELF-E and Pol II or DSIF^28^. As we utilized a 25 nt nascent RNA for our PEC assembly, and as nascent RNAs were reported to stably bind the NELF-E RRM domain beyond a length of 60 nt^30^, the observed RNA-protein crosslinks very likely represent eRNA-protein contacts. When we further analyzed RNA crosslinks to NELF-A, we found them to overwhelmingly lie within the NELF-A tentacle (residues R202, L248, L257, V265, A271, L431, L432, R435, A440, F445, A448, L460), a highly flexible, to date unresolved part of NELF-A that is required for pause stabilization (Fig. 4c)^28^. We found a remarkable spatial correlation between some of the aforementioned RNA crosslinks to the NELF-A tentacle and NELF-A residues K200, K207, K215, K219, K255 and K276, for which protein-protein crosslinking data had established the location of the NELF-A tentacle in context of the entire PEC (Fig. 4d)^28^. Moreover, we found RNA crosslinks to RPB1 (I714), the protrusion domain of RPB2 (S94, L124, D127, L156, F422, G426) and RPB3 (D141, Q157) to all fall within a region that previous protein-protein crosslinking data suggested as the path of the NELF-A tentacle on the surface of the PEC (Fig. 4d) (Vos et al. 2018). Last, we detected multiple RNA crosslinks to RPB1 (I714, N731, L760, N765, E927, L1158, C1159, L1216, R1218) and RPB8 (M145, K146, K147, L148, F150) that are located at the interface between of NELF-AC lobe and Pol II in the PEC structure (Supplementary Fig. 4e). Taken together, the presented protein-RNA crosslinking data allow us to rationalize the length dependency of eRNA-driven NELF dissociation (Fig. 2b-d). Our biochemical data (Fig. 2b-d) argue that a single eRNA molecule has to bind to several RNA binding sites on NELF simultaneously in order to trigger the release of NELF from the PEC. As we detected eRNA binding to both the NELF-A and the NELF-E tentacle (Fig. 4b,c and Supplementary Table 4) – in areas that directly contact Pol II^28^ – we envision that eRNA binding to NELF disrupts critical contacts between NELF and Pol II and leads to its dissociation. In support of the crucial nature of eRNA length for triggering NELF release, we computed a 3D model of *Arc* eRNA (1-200) using the Rosetta-based FARFAR2 tool^42,43^ and using our experimentally determined secondary structure as an input. Intriguingly, *Arc* eRNA (1-200) folds into a structure with dimensions that would render simultaneous eRNA binding to NELF-A/C and NELF-E possible (Fig. 4e). Last, we note that we also observed RNA crosslinks to the NELF-BE lobe (Supplementary Table 4). This suggests that eRNA binding to NELF-BE may be functionally relevant *in vivo*. However, *in vitro* RNA binding to NELF-BE is not capable of triggering NELF release, as our data using the NELF double mutant demonstrate (Fig. 3c-e).

**Fig. 4.**
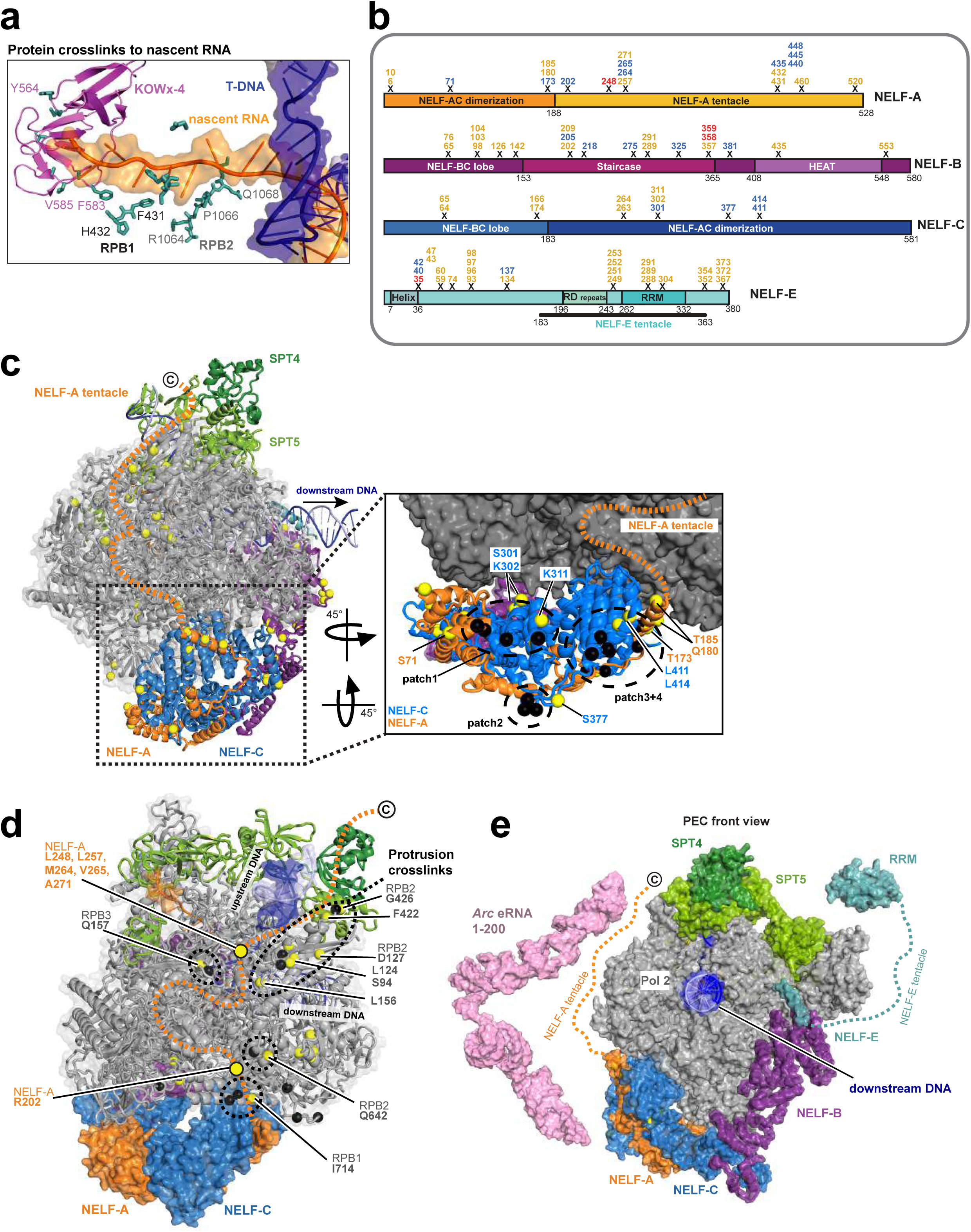
In context of the paused elongation complex, eRNAs bind to positive patches on the NELF-AC lobe, the NELF-A tentacle and NELF-E. **a,** The most frequent protein-RNA crosslinks involving RPB1, RPB2 and the KOWx-4 domain of DSIF can be attributed to nascent RNA. Crosslinks are mapped on the structure of the PEC (PDB code 6GML). Crosslinked residues are shown in stick representation. **b,** RNA crosslinks with an xQuest score > 25 that map to any of the NELF subunits. Crosslinked residues to *Arc* eRNA (1-200) are shown in blue, crosslinks to *Nr4a1*-(b) eRNA (1-100) are shown in orange. Crosslinks found in both experiments are marked in red. To aid interpretation, crosslinked residues that lie within +/− 15 amino acids were visualized as clusters. The domain organization of individual NELF subunits is depicted. **c,** Visualization of RNA crosslinks on NELF-AC, NELF-B and on the Pol II surface. Crosslinked residues are depicted in yellow. The close-up view details RNA crosslinks to the NELF-AC lobe, which are colored in yellow. **d,** Detailed view of RNA crosslinks to the NELF-A tentacle and to the surface of Pol II. The position of the unresolved NELF-A tentacle is indicated as a dashed line as before28. RNA-crosslinked residues on the NELF-A tentacle and on Pol II are shown as yellow spheres. Previously identified protein-protein crosslinks between the NELF-A tentacle and Pol II are shown as black spheres28. **e,** Front view of the PEC (PDB codes 6GML and 2JX2). Alongside the PEC, a 3D model of *Arc* eRNA (1-200) is shown. The Arc eRNA (1-200) model was generated using SHAPE restrained Rosetta modelling with FARFAR43. The model supports that eRNAs need to be 200 nt long in order to efficiently trigger NELF release from the PEC (see Fig. 2,3).

### NELF directly binds to nascent eRNAs *in vivo*

Previous EMSA experiments^30,44^, as well as our own *in vitro* data on PEC assembly (Fig. 4a and Supplementary Fig. 2a), showed that a stable association of DSIF and NELF with Pol II requires synthesis of short nascent pre-mRNAs. This suggests that the interaction between the pausing factors and nascent RNAs are critical for mediating Pol II pausing. However, a direct demonstration of such interaction in an *in vivo* context has never been provided. Moreover, we also wanted to test the *in vivo* relevance of the eRNA-mediated NELF release that we could observe *in vitro* (Fig. 2,3). To this end, we performed eCLIP-seq^45^ in primary neuron culture before and after KCl stimulation. We determined the genome-wide interaction map between NELF-E and RNA during the early stage of activity-induced transcription (Fig. 5a and Supplementary Fig. 5a,b). By nature of eCLIP-seq library preparation, we could estimate the NELF-E crosslinking sites from each sequenced read, which is the first nucleotide of the R2 read (see Methods). To assign the crosslinking sites separately to eRNAs and pre-mRNAs, we utilized the transcript units we had previously defined *de novo* from GRO-seq reads (Supplementary Fig. 5c and Supplementary Table 1). The crosslinking sites were strikingly biased toward the 5’-ends of pre-mRNAs, which is consistent with the previous model that NELF interacts with the 5’-ends of nascent RNAs emerging from transcribing Pol II^30,44^ (Fig. 5b,d,f and Supplementary Fig. 5d,f,h). About 70% of the crosslinking sites were located within the first 200 nt of nascent pre-mRNAs in both unstimulated and KCl-stimulated conditions with a notable enrichment in the first 50 nt. The distribution of the crosslinking sites within eRNAs was similar but more broadly spread to downstream regions (Fig. 5c,e,f and Supplementary Fig. 5e,g,h). Observed bias in the crosslinking sites toward the 5’-ends of nascent transcripts largely correlated with the average NELF-E occupancy, which is highly restricted to the immediately flanking regions of the TSSs (Fig. 5b,c and Supplementary Fig. 5d,e). Given that low levels of NELF binding were detected at some enhancers, it is possible that nascent eRNAs could directly contact the NELF complex bound near the genomic region of its origin. Although our eCLIP-seq analysis does not allow us to distinguish the origin of NELF that interacts with eRNAs, the following features observed from our analysis suggest the possibility of eRNA-dependent disruption of NELF association with paused Pol II at promoters, as previously suggested^17^: 1. There are much lower levels of NELF binding at enhancers than at promoters (Fig. 5g and compare the right y-axis values in Fig. 5b,c); 2. eRNA crosslinking sites show a broader distribution across the length of eRNAs in contrast to pre-mRNA crosslinking sites that are narrowly enriched near the 5’-ends (Fig. 5b-e and Supplementary Fig. 5d-g); 3. When considering pre-mRNAs and eRNAs transcribed from NELF-bound genes (548) and enhancers (144), we detected a notable increase in the crosslinking occurrence within the first 200 nucleotides of eRNAs immediately after KCl stimulation (see discussion) (Fig. 5h-j and Supplementary Fig. 5i-k). Taken together, our eCLIP-seq analysis demonstrates that eRNAs directly interact with NELF in primary neurons, providing the *in vivo* relevance of our *in vitro* findings.

**Fig. 5.**
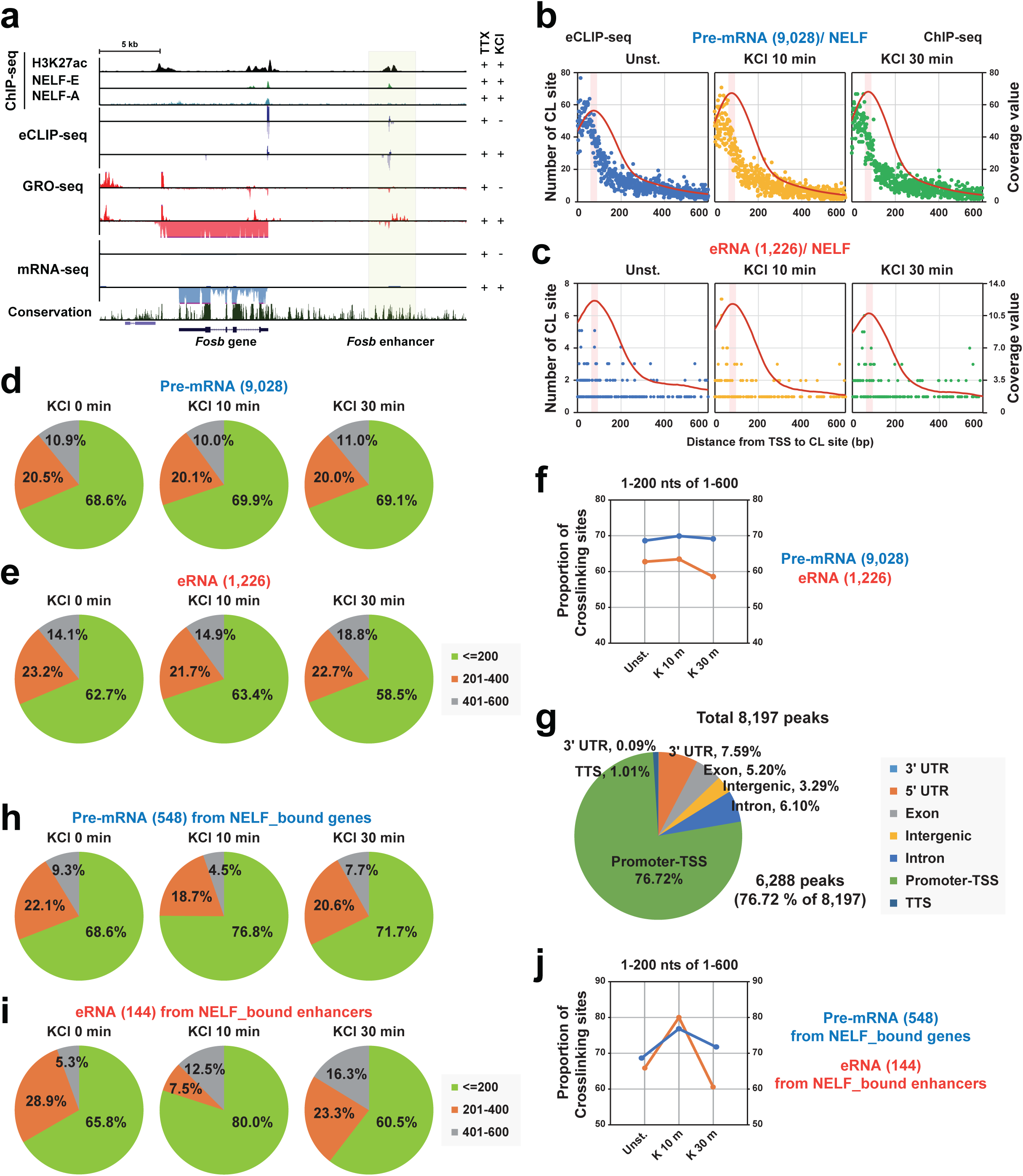
NELF directly binds to nascent eRNAs *in vivo*. **a,** Example track showing one of the activity-induced genes, *Fosb* and its nearby enhancer (shaded area) along with ChIP- (H3K27ac, NELF-A and NELF-E), eCLIP-, GRO-, and mRNA-seq data. (**b** and **c**) The distribution of the crosslinking sites from eCLIP-seq data and NELF ChIP-seq coverage profiles for total annotated pre-mRNAs (9,028) **b,** and total intergenic eRNAs (1,226) (**c**) at different time points after KCl stimulation (0, 10, and 30 min). The number of the NELF crosslinking sites present up to 600 nucleotides from the TSSs as defined by ENCODE annotation of pre-mRNAs (**b**), and up to 600 nucleotides downstream from the *de novo* defined 5’ ends (TSS) of eRNAs (**a**) are shown. Each dot indicates the location of a crosslinking site. Blue, yellow, and green indicates 0, 10, and 30 min KCl stimulation, respectively. The pink shaded area indicates the ChIP-seq peak summit position. (**d**, **e**, **h**, and **i**) Pie charts that show the proportion of crosslinking sites present in three distance windows (1-200, 201-400, or 401-600 nucleotides) for the total annotated (9,028) (**e**) or KCl-up/ NELF_high bound (216) (**h**) pre-mRNAs and the total intergenic (1,226) (**e**) or KCl-induced/ NELF_bound (144) (**I**) eRNAs at different time points after KCl stimulation. See also Supplementary Fig. 7b. (**f** and **j**) Proportion of crosslinking sites within 1-200 nt in the 1-600 nt downstream regions of total pre-mRNAs (9,028) or eRNAs (1,226) (**f**), and of NELF bound pre-mRNAs (548) or eRNAs (144) (**j**). **g,** ChIP-seq analysis of the NELF-E protein. The pie chart shows the genomic distribution of annotated NELF-E peaks.

### eRNA-driven NELF release commands a more efficient release of Pol II from the paused state *in vitro*

NELF release is a hallmark of the transition from promoter-proximal pausing to transcription elongation. As we found eRNA-driven NELF release *in vitro* to resemble pause release *in vivo* (Fig. 2), and as we observed eRNA-NELF interactions *in vivo* (Fig. 5), we next asked whether eRNAs indeed trigger a more efficient release of Pol II from the paused state. To address this question, we established a mammalian *in vitro* pause release assay (see Methods)^28,30,46^ (Fig. 6a). Before examining the effect of eRNAs, we verified that we indeed observe pause stabilization through the presence of both DSIF and NELF using our assay setup (Supplementary Fig. 6a). As expected, by adding *Nr4a1*-(a) or *Arc* eRNA fragments we observed a more efficient release of Pol II from the paused state (Fig. 6b and Supplementary Fig. 6b, upper gel). In striking similarity to our EMSA results (Fig. 2b,c), this effect was critically dependent on eRNA length and less so on the “structuredness” of the tested eRNA (Supplementary Fig. 6b, lower gel). Only *Nr4a1*-(a) eRNA variants with a length of 100 nt or exceeding this length showed increased Pol II pause release. Next, we measured eRNA-induced Pol II pause release using a PEC variant that comprised the NELF-C patch mutant and utilizing both *Arc* and *Nr4a1*-(a) and -(b) eRNAs. As shown in Fig. 6c and Supplementary Fig. 6c, we observed hardly any Pol II pause release with this NELF mutant using *Arc* eRNA (1-50) or (1-100) and substantially diminished rates of release using longer eRNA variants. This is, again, in good agreement with our EMSA data (Fig. 3b and Supplementary Fig. 3b). We note here that the NELF-C patch mutant leads to an overall increase in Pol II pause stabilization (compare quantification in Fig. 6b,c). When using the NELF double mutant (both lacking the NELF-E RRM domain and comprising the NELF-C patch mutants, see Fig. 3c-e), we found Pol II pause release rates similar to those for the NELF-C patch mutant (Fig. 6d and Supplementary Fig. 6d). Taken together, our *in vitro* data on Pol II pause release confirm our prior findings and they further establish that, indeed, eRNAs of sufficient length are able to release Pol II from its paused state by stimulating NELF release. We note that the eRNA-induced increase in Pol II pause release efficiency was not as dramatic as the highly efficient release of NELF following eRNA addition in our EMSA assays (Fig. 2). This is likely the case because NELF release may not automatically jump-start transcription *in vitro*, even more so as NELF induces a tilted conformation of the DNA-RNA hybrid in the active site of Pol II^28^. Moreover, our assay system is not able to recapitulate P-TEFb phosphorylation^33,47^, which may also hinder the efficient resumption of transcription elongation *in vitro*

**Fig. 6.**
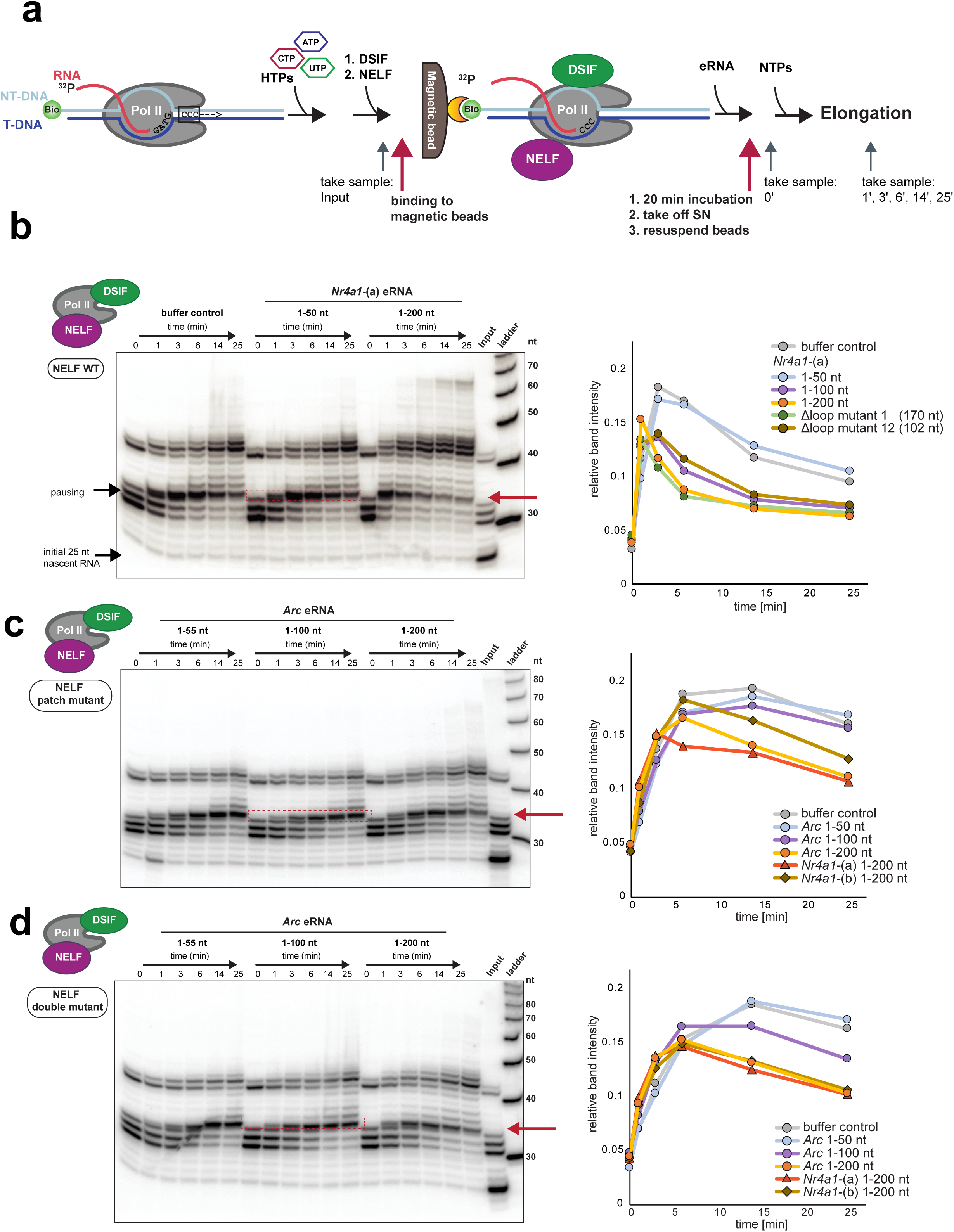
eRNAs facilitate pause release in a length-dependent manner. **a,** Schematic of the mammalian pause release assay. Briefly, a transcription-competent complex that consists of Pol II and a transcription bubble is assembled (see Fig. 2a). As the template DNA contains a G-less cassette, pausing can be induced by GTP omission. DSIF and NELF are then added to reconstitute the PEC and to stabilize the pause. The PEC is then isolated by binding the biotinylated DNA non-template strand to magnetic streptavidin-coated beads. Next, eRNAs are added, the supernatant (SN) is removed, and transcription is allowed to resume by addition of NTPs. **b,** Pause release assay using wild type NELF and *Nr4a1*-(a) eRNA fragments (1-50 and 1-200). The left panel shows the urea PAGE analysis for the *Nr4a1*-(a) 1-50 and 1-200 fragments. Urea PAGE gels for *Nr4a1*-(a) eRNA (1-100) and two Nr4a1-(a) mutants (102 nt - Δloop 12 mutant; 170 nt - Δloop 1 mutant) are shown in Supplementary Fig. 5b. Samples for all experiments were taken just before NTP addition (0 min) and at different time points after NTP addition (1, 3, 6, 14 and 25 min). The “input” sample contains the PEC sample before its affinity purification using the streptavidin-coated beads. Thus, it allows for the visualization of unbound nascent RNA. For the quantification of pause release efficiency (shown on the right) the intensity of the first transcript elongation band past the pause (boxed in red) was analyzed. **c,** Pause release assay using the NELF patch C mutant and *Arc* eRNA (1-55, 1-100 and 1-200). The left panel depicts the urea PAGE analysis for the *Arc* eRNA fragment (1-55), (1-100) and (1-200). Additional data for *Nr4a1*-(a) and *Nr4a1*-(b) 1-200 are shown in Supplementary Fig. 5c. The right panel shows the quantification of all experiments, as described for (b). **d,** Pause release assay using the NELF double mutant. Performed under the same conditions as described in (c).

### NELF binding levels correlate with activity-induced transcription elongation

Because our *in vitro* data revealed that eRNAs abrogate Pol II pausing by releasing NELF from the PEC (Fig. 6), we asked whether this mechanism plays a general role in activity-dependent transcription in neurons. To that end, we further analyzed our GRO-seq profiles as a proxy for nascent Pol II transcription (Fig. 7a-d and Supplementary Fig. 7a-d). We grouped all mRNA genes based on the binding level of NELF-E at the promoter and/or transcriptional inducibility in response to KCl stimulation (mRNA-seq signal FC > 1.5 any time after KCl stimulation) (Fig. 7b and Supplementary Fig. 7b). A total of 623 activity-induced genes (KCl-depolarization induced [KCl-up]) were sub-grouped as NELF-E bound (548) and unbound (75). KCl-up/ NELF-E bound (548) genes were further divided into three different groups based on the level of NELF-E binding (216 NELF-E high, 133 mid, and 199 low) (Fig. 7c and Supplementary Fig. 7b,c). The 252 activity-induced (KCl-up) eRNAs (FC > 1.5) that we found amongst our list of 1,226 intergenic eRNAs (see Supplementary Table 1) were further divided into NELF-E bound (144) and unbound (108) enhancer groups (Fig. 7b,d and Supplementary Fig. 7d). Activity-induced genes show a strong positive correlation between NELF binding levels and transcriptional induction (Fig. 7b,c and Supplementary Fig. 7b,c). In case of activity-dependent enhancers, NELF-bound enhancers show a faster eRNA induction kinetics than those enhancers with no noticeable NELF binding (Fig. 7b,d). We also noticed that eRNAs transcribed from NELF-bound enhancers exhibited a more rapid and transient induction than pre-mRNAs from KCl-up/NELF-E bound genes, which is consistent with previous studies^17,48^ (Fig. 7b). These results collectively suggest that NELF is a functional component in the activity-dependent transcription program. To further examine the function of NELF in Pol II pausing and elongation, we calculated the Pausing Index (PI) based on GRO-seq signals, which can estimate the relative density (and therefore transcription activity) of Pol II at the initiation and elongation stages^49^ (Supplementary Fig. 7e). Activity-induced genes with high levels of NELF binding (KCl-up/NELF_high) showed a decrease in PI at 30 and 60 min after KCl stimulation, indicating an increase in Pol II elongation (Figures 7E and S7F). However, such elongation stimulatory effect was not observed in activity-induced genes with weak or no NELF binding. There, a NELF-dependent increase in eRNA elongation was also observed but to a much lesser degree (Fig. 7f and Supplementary Fig. 7g). Taken together, these results suggest that in primary neuron culture NELF is part of the activity-dependent gene expression program through elongation control. NELF might also control eRNA transcription as about half of the activity-induced enhancers are bound by NELF, albeit at much lower levels than the activity-induced promoters.

**Fig. 7.**
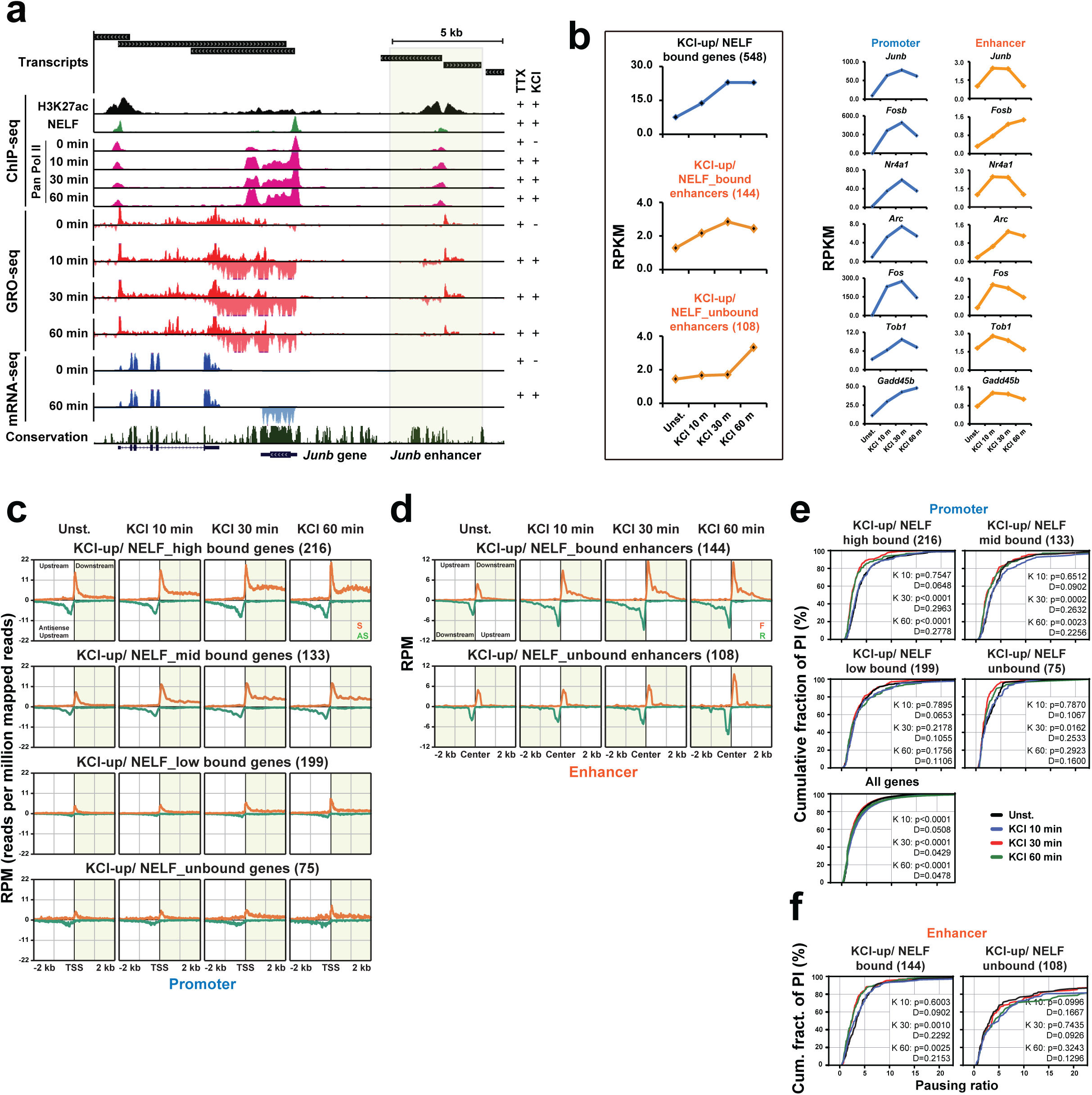
NELF occupancy is correlated with Pol II pause release and causal to transcription induction *in* vivo. **a,** Example track that shows an activity-induced gene, *Junb*, and the nearby enhancer (shaded area). Transcript units determined by *de novo* transcript calling are shown on the top together with tracks for ChIP-(H3K27ac, NELF-E, and Pan Pol II), GRO-, and mRNA-seq. **b,** Transcription induction profiles (RPKM values) of pre-mRNAs and eRNAs, determined based on GRO-seq data in response to 0, 10, 30, and 60 min KCl treatments. See also Supplementary Fig. 7b. There are a total of 623 activity-induced genes (determined by mRNA-seq), which is then divided into groups of NELF-E bound (548) and unbound (75) genes. There are a total of 1,226 intergenic enhancers that transcribe eRNAs. Out of these, activity-induced enhancers are determined based on eRNA induction (> 1.5-fold change at any time point after KCl stimulation), which is then further divided into NELF-E bound (144) and unbound (108). Exemplary induction profiles of genes and their corresponding eRNAs are shown on the right. (**c** and **d**) The average GRO-seq profiles of three different gene groups based on the levels of NELF binding (216 NELF high, 133 mid, 199 low or 75 unbound) (**c**) and KCl-induced/ NELF_bound (144) or unbound (108) enhancers (**d**). The profiles are shown for ±2 kb regions centered on the TSSs or the enhancer centers upon 0, 10, 30 and 60 min KCl stimulation. Orange lines denote the sense strand at promoters and the forward strand at enhancers. Green lines denote the anti-sense strand at promoters and the reverse strand at enhancers. (**e** and **f**) Pausing index (PI) profiles of four different gene groups based on their levels of NELF binding with all genes (**e**) and KCl-up/ NELF_bound (144) or unbound (108) enhancers (**f**). Each gene group is analyzed at 4 time points (unstimulated (Un), 10, 30 and 60 min KCl). The PI is defined as in Supplementary Fig. 7e. Statistical significance between cumulative probability graphs was determined by the Kolmogorov-Smirnov test.

## DISCUSSION

### To detach NELF, eRNAs must bind to multiple sites on the paused elongation complex simultaneously

Our mechanistic study directly links eRNAs and mammalian Pol II for the first time and it fundamentally advances our previous findings^17^. What is more, in addition to the well-characterized NELF-E RRM domain^36–38^, our data also shed light on the role of previously reported additional nucleic acid binding interfaces on NELF^40^. We find these binding sites – in particular on the NELF-AC lobe - to be critically involved in eRNA binding (Fig. 3,4). Moreover, the striking length dependency of eRNA-driven NELF dissociation (Fig. 2) and the significant distance between the NELF-AC lobe and the NELF-E RRM domain, both of which we find directly involved in promoting NELF dissociation upon eRNA binding (Fig. 3), allow us to suggest that eRNAs simultaneously occupy several binding sites across the PEC to trigger NELF release (Supplementary Fig. 8). Furthermore, we envision that eRNAs are attracted to the PEC in a sequence-independent manner by the positively charged patches on NELF-AC. This binding event could then spawn further interactions between the eRNA and the NELF-A tentacle and the distant NELF-E RRM domain. Finally, the sum of all eRNA-NELF interactions then triggers NELF release. In support, the NELF-A tentacle was hypothesized to significantly contribute to the overall affinity of the NELF complex for RNA^40^, and we found multiple protein-RNA crosslinks at the interface of NELF-AC and Pol II (Supplementary Fig. 4e)^40^.

### Unpaired guanosines in enhancer RNAs and transcriptional condensates go hand in hand

Against initial expectations eRNAs neither share common structural motifs, nor do they possess sequence motifs that determine their function (Fig. 1,2). In contrast, we find that enhancer RNAs must only meet two loose criteria to be able to efficiently abrogate Pol II pausing by detaching NELF from Pol II: 1. They need to be longer than 200 nt, and 2. multiple unpaired guanosines need to be distributed along the entire enhancer RNA (Fig. 2). The preference of NELF for unpaired guanosines connects well to previous findings. First, the NELF-AC subcomplex was reported to bind single-stranded RNA with a GC content of 60%, but not RNA with a GC content of 44%^40^. A closer look at the utilized RNA sequences revealed, however, that, while the RNA representing a GC content of 60% comprised stretches of guanosines (14 Gs in total), the RNA representing a GC content of 44% contained no guanosines at all. Second, preferential binding of NELF to guanosines was also reported for the *D. melanogaster* homolog of NELF-E and its isolated RRM domain^39^. The study identified the sequence CUGGAGA as NELF binding element (NBE). Mutating all guanosines to adenosines within the NBE abolished *D. melanogaster* NELF-E RNA binding, underscoring the preference of NELF for guanosines. Interestingly, a comparison of the sequences of fly and human NELF-E reveals that the *D. melanogaster* homolog is lacking the RD repeat domain^39^. This fact might explain why human NELF-E binds RNA in a less sequence-specific manner and allows for binding of the broad range of RNAs that our data reveal.

A lack of well-defined motifs that render eRNAs active is not without mechanistic precedent. PRC2 (Polycomb repressive complex 2), a histone methyltransferase that establishes repressed chromatin by trimethylating histone H3 at lysine 27, overall shows promiscuous RNA binding^50,51^. However, the complex preferentially binds to unstructured G-rich sequences and to G-quadruplexes^52^. This is reminiscent of our RNA binding data on NELF (Fig. 2-4), with the notable exception that we find no evidence for a role of G-quadruplexes in eRNA function. Intriguingly, G-tract-containing RNAs sequester PRC2 from nucleosome substrates and thereby lead to gene activation^53,54^. Hence, decoying protein factors with overall promiscuous RNA binding activity by high-affinity binding to G-rich RNAs might be a general concept in RNA-dependent regulation of gene expression. Last, a lack of specificity of RNA-protein interactions might also be compensated for by compartmentalization into sub-organellar structures. This is exactly what was proposed for the regulation of eukaryotic transcription by transcriptional condensates^55,56^. As RNA binding increases the phase separation properties of RNA-binding proteins^57,58^, we envision that eRNAs contribute to the formation of transcription condensates. They likely do so by significantly increasing the valency of the network of interactions that lies at the heart of phase-separated foci^56^. Moreover, we hypothesize that the presence of enhancer RNAs in the same transcriptional condensate as Pol II and other pausing factors will greatly enhance their capacity to regulate gene expression, *e.g.* by abrogating promoter-proximal pausing. Indeed, a recent *in vitro* study under physiologically relevant conditions demonstrated that eRNAs transcribed from super-enhancers influence condensate formation by purified Mediator complex^59^.

### eRNAs can directly interact with NELF in an *in vivo* context

Our eCLIP-seq analysis in primary neuron culture demonstrates that NELF directly interacts with both pre-mRNAs and eRNAs with a strong bias toward their 5’-ends. This provides the first genome-wide *in vivo* evidence that supports the model that a stable association of NELF with paused Pol II is mediated by its interaction with nascent RNA^30,44^. The observed direct interactions between eRNAs and NELF in primary neurons further suggest that the eRNA-mediated NELF release from the PEC demonstrated by our reconstituted system could be part of a transcriptional induction mechanism that operates in an *in vivo* context. However, we cannot rule out that some eRNA eCLIP-seq signal stems from crosslinking of eRNAs to NELF bound at enhancers, as low levels of NELF binding peaks are also present at enhancers (Fig. 5). With this caveat, the following observations are consistent with the possibility that eRNAs mediate destabilization of NELF at promoters *in vivo*. The distribution of eRNA crosslinking sites is broader than that of pre-mRNAs relative to the average NELF binding peak area (Fig. 5b,c and Supplementary Fig. 5d,e). The pre-mRNA crosslinking sites are tightly enriched in less than ∼50 nucleotide regions from their 5’-ends, which is even more upstream than the NELF peak (∼60-70 bp downstream of the TSSs). In contrast, a significant number of eRNA crosslinking sites are observed to occur throughout up to ∼1,000 nt regions from their 5’-ends, which coincides with the typical size of an eRNA transcription unit^11^. This could indicate that, unlike pre-mRNAs, eRNAs interact with NELF during and even after synthesis of their full-length transcripts, which indirectly suggests a possibility of making contacts with the promoter-bound NELF. The broad distribution of crosslinking sites also fits nicely with the length requirement of eRNAs for efficient NELF release proposed by our *in vitro* data (Fig. 2,4). Another feature of eRNAs compatible with our model is their fast induction kinetics. For several activity-induced genes and enhancers, we observed that eRNA induction peaks earlier than pre-mRNAs from their target genes, a finding also observed by others in various enhancer-gene contexts (Fig. 7b and Supplementary Fig. 7b)^4,17,48, 60–63^. Therefore, whether or not eRNA production is under the control of NELF activity, a population of eRNAs that are fully transcribed before pre-mRNA production rises can be available for interaction with promoter-bound NELF. In this regard, it is worth pointing out that a previous study in *Drosophila* also suggested that Pol II at enhancers undergoes more rapid pause release than at promoter regions^10^. This model is well suited with a broader distribution of the crosslinking sites across the length of eRNAs - as mentioned above - and also with the observation that the eRNA crosslinking frequency is prominently increased at 10 min after KCl stimulation than unstimulated and 30 min KCl conditions (Fig. 5h-j and Supplementary Fig. 5i-k). We can further infer that eRNA-mediated NELF detachment from paused Pol II is more feasible in a transcriptional condensate environment created by enhancer-promoter loops, where eRNAs are closely confined with promoter-associated transcription factors and Pol II.

## METHODS

### Mouse cortical neuron culture and stimulation

Mouse cortical neurons were dissected at embryonic day 16.5 (E16.5) and cultured in Neurobasal media (NB) (21103-049, ThermoFisher Scientific) supplemented with 2% B-27 (17504-044, ThermoFisher Scientific) and 1% Glutamax (35050, ThermoFisher Scientific). For KCl depolarization, neurons at DIV 7 were made quiescent by 1 μM tetrodotoxin (TTX; 1078, Tocris, Minneapolis, MN) overnight. 55 mM KCl was then added for the indicated length of time.

### Global run-on sequencing (GRO-Seq)

GRO-seq was performed as previously described^18,64^. Briefly, 10 million nuclei per sample were used for global run-on, and base hydrolysis was performed as previously described^65^. Nascent RNA was immunoprecipitated with anti-BrdU antibody-conjugated beads (sc-32323 AC, Santa Cruz Biotech, Santa Cruz, CA). Purified run-on RNA was subjected to polyA tailing by Poly(A)-polymerase (14.06 U; M0276L, NEB, Ipswich, MA) for 12 min at 37 °C. PolyA-tailed RNA was subjected to another round of immunopurification by using anti-BrdU antibody conjugated beads. Reverse transcription was then performed using SuperScript III Reverse Transcriptase (200 U; 18080-044, ThermoFisher Scientific) with RT primer (pGATCGTCGGACTGTAGAACTCT/idSp/CCTTGGCACCCGAGAATTCCATTTTTTTTTTTTTTTTTT-TTVN) for 2 hr at 48 °C. Extra RT primers were removed by Exonuclease I (100 U; M0293S, NEB, Ipswich, MA) for 2 hr at 37 °C. cDNAs were fragmented with basic hydrolysis and size selected (130–500 nucleotides) in a 6-8 % polyacrylamide TBE-urea gel. Purified cDNAs were circularized using CircLigase (50 U; CL4111K, Epicentre) for 2 hr at 60 °C and relinearized at the basic dSpacer furan with Ape 1 (15 U; M0282S, NEB, Ipswich, MA) for 2 hr at 37 °C. The relinearized single-stranded DNA template was subjected to PCR amplification by using barcoded primers for Illumina TrueSeq small RNA sample and Phusion High-Fidelity DNA Polymerase (2 U; M0530L, NEB, Ipswich, MA). Subsequently, PCR products were size-selected in 6 % polyacrylamide TBE gel (175∼400 bp) and purified. The final libraries were sequenced using Illumina NEXTSEQ 500 following the manufacturer’s instructions.

For analysis, low quality and adapter sequence of the raw FASTQ reads were trimmed using Trim_galore with default parameters. To trim polyA sequence, we used Trim_Galore again with ‘--poly’ parameter. The trimmed reads were aligned to the mm10 GENCODE annotation using STAR. Transcripts were called using ‘findPeaks’ of HOMER package with ‘--minReadDepth 200’ parameter. Samtools and the HOMER package were used to make visualization tracks and RPKM calculations. To calculate RNA expression values by gene accurately, we used Homer package. To generate GRO-seq coverage plots, we used HOMER. For HOMER to draw promoter regions, “makeTagDirectory” program was used. Before making tag directories, we converted BAM files to BED files using ‘bedtools bamtobed’. Using HOMER’s in-built Perl scripts, annotatePeaks.pl data were used to create coverage plots. Pausing indices were calculated as shown in Supplementary Fig. 7e using HOMER’s in-built perl script, ‘analyzeRepeats.pl’. The widths of intervals used to calculate pausing indices were determined from analysis of GRO-seq alignments. The promoter-proximal region was defined as 100 bp upstream to 200 bp downstream of the TSSs. The gene body was defined from 400 to 800 bp downstream of the TSSs. In each window, GRO-seq read density was calculated.

### Identification of enhancers

Enhancers were identified according to MACS-called H3K27ac-enriched peaks. Firstly, H3K27ac peaks were identified in KCl-stimulated neurons (GSM1467417 and GSM1467419)^19^ with their corresponding input controls (GSM1264367 and GSM1467416, respectively) using MACS. Mapped reads of two biological replicates in each condition group were merged before the enhancer calling analysis using Bamtools. Then, the ranking of super enhancer (ROSE) algorithm was used to define super-enhancers with the identified H3K27ac peaks. The H3K27ac peaks that were not overlapped with the super-enhancers (SEs) or promoter regions of known genes were defined as typical-enhancers (TEs). We used the pool of SEs and TEs as total enhancers in this study. To identify *de novo* enhancer transcripts from total GRO-seq transcripts, H3K27ac peaks within ±2 kb region from TSSs and gene body regions were removed from total enhancers. Then, total GRO-seq transcripts overlapped with total enhancers above were defined as *de novo* enhancer transcripts.

### mRNA-seq

mRNA-seq library was constructed using TruSeq RNA Library Preparation Kit (Illumina) according to the manufacturer’s instructions. FASTQ reads from Genomics Core at UTSW were mapped to UCSC’s mm10 genome using Tophat with options “-a 8 -m 0 -I 500000 -p 8 -g 20 --library-type fr-firststrand --no-novel-indels --segment-mismatches 2”. Since this data was strand-specific, we used the “-library-type fr-firststrand” option of Tophat. Reads with low mapping quality (<10) were removed using SAMtools. Duplicate reads were marked by Picard MarkDuplicates (https://broadinstitute.github.io/picard/). Tag directories for each sample were created using “makeTagDirectory” program. RNA expression was quantified using HOMER’s inbuilt Perl script “analyzeRepeats.pl.” These scripts offer flexibility to calculate expression values as reads per kilobase per million mapped reads (RPKM) normalized to 10 million at introns, exons and genebody locations. “makeUCSCfile” from HOMER was used to create bedGraph files at 1bp resolution and created bigWig files for visualization on UCSC genome browser. All coverage values were normalized to 10 million reads. We set expressed gene criteria as “RPKM values higher than 1 at least in one of 6 samples (3 conditions, 2 replicates for each condition)”. The subsequent RNA-seq analyses were performed with these 12,739 genes. To call significant differentially expressed genes (DEGs), we set our criteria as “fold change of RPKM is more than 1.5 and FDR of DESeq2 is less than 0.05 in both replicates”.

### Depletion of ribosomal RNA and SRP-RNA from total RNA isolated from mouse cortical neurons

To deplete rRNA from total RNA isolated from mouse cortical neurons a commercially available kit (RiboCop rRNA depletion kit V1.2, Lexogen) was used according to the manufacturer’s instructions with some modifications. Briefly, for the simultaneous, additional depletion of the signal-recognition particle RNA (SRP-RNA) we utilized four biotinylated DNA oligos (see below) and mixed those with the probe mix (PM) directed against rRNA that is part of the Lexogen kit. 500 ng of total RNA from unstimulated or stimulated (45’ KCl treatment) primary neurons were used for one round of depletion. In total, 115 µL of magnetic streptavidin beads (75 µL depletion beads from the kit + 40 µL Dynabeads MyOne Streptavidin C1, ThermoFisher Scientific) were used. All buffer volumes of the depletion kit were scaled according to the manufacturer’s protocol. The hybridization mix was prepared from total RNA, 6.2 µL hybridization solution (HS), 5 µL probe mix (PM) (volume not scaled) + 3.5 pmol anti-SRP oligo-mix in a final volume of 54 µL. Following the first depletion step, a second round of depletion was performed to ensure that all depletion oligos were removed. To that end, the supernatant containing the rRNA- and SRP-RNA depleted RNA was transferred into a fresh reaction tube and supplemented with 30 µL of pre-equilibrated magnetic streptavidin beads. The mix was incubated at room temperature for 10 min and subsequently at 52 °C for 10 min. The cleared supernatant was then transferred to a fresh reaction tube and applied to spin-columns (RNA Clean & Concentrator-5 columns, Zymo-Research) according to the manufacturer’s instructions. For one Exo-seq library sample, RNA from three depletion reactions (3x 500 ng total RNA) were pooled prior to column purification.

anti_SRP_oligos (for SRP-RNA depletion):

#1: 5’-Biotin-TACAGCCCAGAACTCCTGGAC TCAAGCGATCCTCCTG

#2: 5’-Biotin-ATCCCACTACTGATCAGCACG GGAGTTTTGACCTGCTC

#3: 5’-Biotin-TCACCATATTGATGCCGAACT TAGTGCGGACACCCGATC

#4: 5’-Biotin-CTATGTTGCCCAGGCTGGAGT GCAGTGGCTATTCACAG

### Exo-seq library preparation and data analysis

The Exo-seq library construction from two biological replicates of KCl (30 min)-treated and one untreated sample was performed as described^20^, with the following alternations. Instead of poly(A)-selected RNA we started with an rRNA and SRP-RNA depleted sample (prepared from 1.5 µg total RNA input, see before). DNA oligos were altered to include a linker 1 (3‘-adapter: 5Phos/TGGAATTCTCGGGT GCCAAGG/ddC) and a linker 2 (5’-adapter: 5Phos/GATCGTCGGACTGTAGAAC TCTGAAC/ddC) sequence based on the adapters of the Illumina TruSeq small RNA sample prep kit. For the final PCR enrichment step, which was performed for 14 cycles, TruSeq RP1 (forward) and RPI Index (reverse) primers from the TruSeq kit were used. The library was cleaned up and size-selected by two consecutive rounds of binding to 1.2x and 0.8x SPRI beads (Agentcourt AMPure XP beads, Beckman Coulter). The integrity and size distribution of the final libraries (two biological replicates of KCl-treated and one of untreated) were analyzed on a Bioanalyzer before they were applied to a NextSeq 500 platform for a single-end 75 nt sequencing run.

Raw reads were processed using *Cutadapt* for adapter trimming, retaining all reads with a minimum read length of 20 nt after trimming. Remaining rRNA- and tRNA reads were removed using *SortMeRNA* and *Bowtie*, respectively. rRNA and tRNA reference files were obtained from the UCSC table browser. Processed reads were then aligned to the mouse genome (mm10, December 2009) using *STAR*, allowing two mismatches and removing multimappers. 5’-end coverage of mapped reads was calculated for both strands separately using the bedtools genomecov utility with -bg and −5 parameters. TSSCall was then run on the generated bedgraph files with standard parameters for identification of transcription start sites (TSS). We defined extragenic TSSs as TSSs that do not occur within a RefSeq-annotated gene ± 2kb. By using these criteria 129,161 extragenic TSSs were identified for Rep1 and 131,312 for Rep2. Called TSSs were then overlapped with *de novo* GRO-seq defined enhancer transcript units allowing an offset of ±200 nt. A single TSS was then selected for each identified enhancer locus. This selection was based on read coverage and distance to the 5’-end of the respective GRO-seq transcript. This resulted in 977 TSSs for Replicate 1 and 1,039 for Replicate 2. Activity-induced enhancers were defined based on GRO-seq fold-change (FC > 1.5) between stimulated and unstimulated conditions. In a last step, 39 high quality eRNA TSSs were selected for structure mapping. Among these 33 eRNA TSSs were derived from activity-induced eRNAs, as defined by GRO-seq data.

### *In vitro* transcription and purification of enhancer RNAs

eRNAs used for EMSAs, transcriptional pause release assays and for SHAPE-MaP were produced by T7 RNA polymerase-mediated *in vitro* run-off transcription using PCR generated DNA templates (adapted from^66,67^). For this purpose, mouse eRNA sequences were cloned via Gibson assembly into pUC18 vectors containing a T7 promoter sequence (5’-TAATACGACTCACTATAGG) in front of the eRNA sequence. Then, a PCR templates was generated using a T7 promoter-spanning forward primer and an eRNA-specific reverse primer. Depending on the desired yield of each eRNA, 200 µL (for SHAPE-MaP eRNA) or 0.8-1.2 mL (for EMSA) transcription reactions were set up. The transcribed eRNAs were purified by urea-PAGE, passively eluted into RNA buffer (25 mM K-HEPES pH 7.4, 100 mM KCl, 0.1 mM EDTA) by the crush and soak method^68^ and concentrated to a volume of 0.5 – 1 mL using centrifugal filters (Amicon Ultra-4 /-15, Millipore) with a molecular weight cutoff (MWCO) depending on the molecular size of the produced fragment (1-50 nt – 3 kDa; 1-100 nt - 10 kDa; 1-200 nt – 10 kDa or 30 kDa). Subsequently, eRNAs were subjected to size-exclusion chromatography using RNA buffer as running buffer on a Superdex 200 Increase 10/300 GL column to purify RNA monomers from aggregates. After elution from the column, the monodisperse peak fractions were pooled and concentrated to eRNA concentrations in the range of 10 – 20 µM.

### SHAPE-MaP library preparation and data analysis

39 eRNAs (1-200 nt) for SHAPE-MaP were produced as described above, except that eRNA fragments for SHAPE-MaP were flanked with a 20 nt long 5’-linker (5’-GGC CAT CTT CGG ATG GCC AA) and 43 nt long 3’-linker (5’-TCG ATC CGG TTC GCC GGA TCC AAA TCG GGC TTC GGT CCG GTT C)^69^. Prior to chemical probing, each purified eRNA (10 pmol in 18 µL) was incubated in 1x HEMK buffer (50 mM K-HEPES pH 7.0, 0.1 mM Na-EDTA, 150 mM KCl, and 15 mM MgCl_2_) for folding at 37 °C for 30 min (adapted from^70^). Chemical probing and library preparation were carried out according to the small RNA workflow^25^. The folded RNA sample was split into two samples: each sample was either treated with 1 µL of 100% DMSO (DMSO control sample) or 1 µL of 100 mM 1-methyl-7-nitroisatoic anhydride (1M7 modified sample) at 37 °C for 5 min. The treatment was repeated for a second round of modification to achieve higher modification rates. The volume of the modified RNA was adjusted to 30 µL with DEPC-treated water, the RNA was cleaned up with 1.8x AMPure XP beads (Beckman Coulter) according to the manufacturer’s instructions and eluted in 15-25 µL of DEPC treated water. 10 µL of the purified RNA sample were subjected to reverse transcription using a specific SHAPE-MaP RT primer complementary to the 3’ linker region, as described^25^. For eRNAs which yielded no DNA product the temperature during reverse transcription was increased to 50 °C. The cDNA was cleaned up with 1.8x AMPure XP beads and eluted in 35 µL DEPC-treated water. Subsequently, the first and second PCR steps were performed as follows: Two distinct Index-primers were used for the DMSO control and the 1M7-modified libraries. The DNA from the 1^st^ PCR was purified with 1.0x AMPure XP beads and from the 2^nd^ with 0.8x. Final libraries were eluted in 25 µL. Each sample was analyzed on a Fragment Analyzer (Agilent) and concentrations were fluorometrically quantified using Qubit (ThermoFisher Scientific). All individual samples were diluted with library dilution buffer (10 mM Tris-HCl pH 8.0, 0,1% (v/v) Tween20) to 5 nM and pooled equimolar. Libraries were sequenced on a NextSeq 500 platform with a Mid-Output Next-Seq kit in a 150 bp paired-end mode and approximately 30% PhiX due to the low complexity of the library mix.

SHAPE-MaP sequencing reads were analyzed by the *ShapeMapper 2* tool. The software was executed with the default parameters, which align the reads to the given eRNA reference sequences using Bowtie 2 (default option was used) and compute the read-depth of each eRNA along with the mutation rates and SHAPE reactivities. eRNA secondary structures were predicted with *RNAStructure*^27^ using the SHAPE reactivities per nucleotide as constraints. The *MaxExpect* structure (maximum expected accuracy structure)^27^, was selected for visualization of the secondary structure using the *StructureEditor* (belongs to RNAStructure software package). Base pairing probability files (.dp) were generated from *MaxExpect* structure .pfs files (partition save file) by *ProbabilityPlot* with the -t option for text file output. For each base we selected the base pairing with maximum probability by utilizing a custom R script. Bases with pairing probabilities > 0.9 were defined as structured bases. Using those base pairs as a measurement for the “structuredness” of eRNAs, we compared the percentage of structured bases over the entire set of 39 eRNAs. Sequences of eRNAs were extracted from position 1 to 1000 using gffread. Sequences were then binned in non-overlapping bins of 200 nucleotides and analyzed for their sequence content. Differences in base content were tested for significance with pairwise t-test using a custom R script.

### Expression and purification of proteins for *in vitro* experiments

#### Purification of mammalian RNA polymerase II

*Sus scrofa* Pol II was purified as described with minor alterations as follows^29,71^. Pig thymus was collected from the Bayerische Landesanstalt für Landwirtschaft (LfL) in Poing, Germany. The bed volume of the 8WG16 (αRPB1 CTD) antibody-coupled sepharose column was 3 mL. The final SEC step was omitted. Instead, Pol II peak fractions after anion-exchange chromatography (UNO Q, Bio-Rad) were pooled before buffer exchange to Pol II buffer (25 mM Na-HEPES pH 7.5, 150 mM NaCl, 10 µM ZnCl_2_,10% (v/v) glycerol, 2 mM DTT) 100 kDa Amicon centrifugal filters (Millipore). Pol II was concentrated to 2.6 mg/mL (5 µM), aliquoted, flash-frozen in liquid nitrogen and stored at −80°C.

#### Expression of NELF variants and DSIF

Bacmids for protein expression were made in DH10EMBacY cells^72^. For the generation of V_0_ virus, bacmids were transfected into SF21 insect cells in SF-4 Baculo Express medium (Bioconcept) at a density of 0.8×10^6^ cells/mL. YFP fluorescence was monitored as a proxy for transfection efficiency and cells were kept at a density of 0.8×10^6^ cells/mL until 100% of cells showed YFP fluorescence. The supernatant, containing the V_0_ virus, was subsequently isolated and utilized to infect High Five cells at a 1:10 through 1:20 ratio. For protein expression High Five cells were kept at a density of 1.0-1.2×10^6^ cells/mL. Protein expression was allowed to proceed for 48 h before cells were harvested by centrifugation. Cell pellets were resuspended in 60 mL (per 1 L expression culture) lysis buffer (for NELF: 20 mM Na-HEPES pH 7.4, 300 mM NaCl, 10% (v/v) glycerol, 30 mM imidazole, 2 mM DTT; for DSIF: 50 mM Tris-HCl pH 8.0, 150 mM NaCl, 2 mM DTT) supplemented with the protease inhibitors pepstatin A (1 µg/mL), leupeptin (1 µg/mL), Benzamidin-hydrochloride (0.2 mM) and PMSF (0.2 mM). Cells were directly used for protein purification or flash frozen in liquid nitrogen and stored at −80 °C.

#### Purification of NELF complex variants

The purification protocol for the wild-type NELF complex and all NELF complex variants (NELF-EΔRRM, NELF patch mutant and NELF double (patch + ΔRRM) mutant) were adapted from a prior protocol^40^. Following Ni-NTA affinity chromatography (Qiagen), the NELF containing protein fractions were pooled, filtered and loaded onto a pre-equilibrated (20 mM Na-HEPES pH 7.4, 300 mM 10 % (v/v) glycerol, 2 mM) Resource Q column (6 mL column volume). As the 4-subunit NELF complex does not bind to the column, the flow through was collected, diluted to 150 mM NaCl, reloaded onto the Resource Q column and eluted with a linear salt gradient to 1 M NaCl (The first Resource Q step allows for the removal of trimeric NELF-ABD complexes). Peak fractions were pooled, concentrated using 30 kDa Amicon concentrators (Millipore) and loaded onto a Superose 6 10/300 column that had been equilibrated in SEC (size exclusion chromatography) buffer (20 mM Na-HEPES pH 7.4, 150 mM NaCl, 2 mM DTT). Peak fractions after SEC were pooled and concentrated to approx. 30 µM (∼7.0 mg/mL). Finally, glycerol was added to 10 % (v/v) before the protein was aliquoted, flash-frozen in liquid N_2_ and stored at −80°C.

#### Purification of DSIF

DSIF carried a C-terminal 2xStrep-tag on SPT5. Cells were lysed by sonication, lysates were then cleared by ultracentrifugation (Type 45 Ti (Beckman), 35 min, 33 000 rpm, 4 °C). Subsequently, the supernatant was applied to 1 mL Strep-Tactin beads (IBA) that had been equilibrated in binding buffer (50 mM Tris-HCl pH 8.0, 150 mM NaCl, 2 mM DTT). Beads were then washed with the same buffer (40 CV) and the protein was eluted with 20 mL (1 mL fractions) elution buffer (binding buffer supplemented with 2.5 mM desthiobiotin). Elution fractions were analyzed by 10% Bis-Tris PAGE, DSIF-containing fractions were pooled and applied to a Resource Q column. DSIF was eluted with a linear salt gradient from 150 mM to 1 M NaCl. Peak fractions were pooled, concentrated using 30 kDa Amicon concentrators (Millipore) and applied to a Superose 6 10/300 column in SEC buffer (20 mM Na-HEPES pH 7.4, 150 mM NaCl, 2 mM DTT). Peak fractions were pooled, concentrated to 7.5 µM (∼1.1 mg/mL), glycerol was added to 10 % (v/v) before the protein was aliquoted, flash frozen in liquid nitrogen and stored at - 80 °C.

### Electrophoretic mobility shift assays (EMSAs)

The PEC was formed on a nucleic acid scaffold using the following oligos: RNA 5’-UCU AGA ACU AUU UUU UCU UAC ACU C (25 nt), template DNA 5’-CTG TAG ACT GAC CAA GTT GTC CCG TAG GAG TGT AAG AGA TAT ATG GTA G (49 nt), and non-template DNA 5’-CTA CCA TAT ATC TCT TAC ACT CCT ACG GGA CAA CTT GGT CAG TCT ACA G (49 nt).

The sequence of the shorter 15 nt RNA, used in control experiments, matched the 15 nucleotides from the 3’-end of the longer 25 nt RNA. Prior to the assay, 100 pmol of RNA were 5’-end labeled with [γ- ^32^P]-ATP (3000 Ci/mmol, *Perkin Elmer*) using T4 polynucleotide kinase (*NEB*) according to the manufacturer’s protocol. The labeled RNA was purified by ethanol precipitation and subsequently annealed to 100 pmol of template DNA in annealing buffer (20 mM Na-HEPES pH 7.4, 100 mM NaCl, 3 mM MgCl_2_, 10% (v/v) glycerol) in a final volume of 20 µL. To this end, the mixture was heated at 95°C for 5 min and cooled down to 20°C in 1°C /min steps in a thermocycler (as previously described by Vos et al., 2018). After annealing, the sample was diluted 1:1 in DEPC-treated water to achieve a final concentration of 2.5 nM hybrid in 40 µL. Subsequently, all amounts refer to one individual EMSA sample loaded onto one gel lane. EMSA samples were typically prepared in a *n*-fold master mix, which was then split into *n* samples before the addition of different eRNAs. Each incubation step, if not explicitly mentioned, was performed at 30 °C for 15 min. The final sample volume was 8 µL. The PEC was formed by incubating pre-annealed RNA-template DNA hybrid (0.8 pmol) with Pol II (1.2 pmol). Then, non-template DNA (1.6 pmol) was added and incubated. Next, the reaction was supplemented with 5x EMSA buffer to achieve a final concentration of 20 mM Na-HEPES pH 7.4, 100 mM NaCl, 25 mM KCl, 3 mM MgCl_2_, 2 mM DTT. Subsequently, DSIF (2.4 pmol) and NELF (1.2 pmol) were added. Each added protein was incubated with the reaction mixture as described above. Finally, eRNAs were added in increasing amounts (1.2, 2.4, 4.8, 7.2, 9.6, 14.4, 19.2 pmol) to yield final concentrations of 0.15, 0.3, 0.6, 0.9, 1.2, 1.8, 2.4 µM. If an EMSA gel contains a series of six instead of seven different eRNA concentrations, then the highest concentration (2.4 µM) was omitted. The samples were incubated for 15-20 min at room temperature, subsequently supplemented with 1.5 µL of EMSA loading dye (20 mM Na-HEPES pH 7.4, 60% (v/v) glycerol) and loaded on a pre-chilled vertical 3.5% native acrylamide gel (0.5x TBE), which had been pre-run for 30 min at 90 V and 4 °C (used pre-chilled 0.5x TBE running buffer and put gel running chamber on ice during electrophoresis). Samples were separated for 1 h and 30 min. Subsequently, gels were dried and exposed to storage phosphor screens for 3 h or overnight, depending on the level of radioactivity. The phosphor screens were read out by a CR 35 image plate reader (Elysia Raytest). Gels were densitometrically analyzed using ImageJ and the Pol II-DSIF fraction was plotted against the eRNA concentrations using Prism 9.

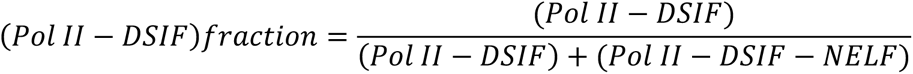

The pseudo-binding curves were fitted with a single site quadratic binding equation as previously described^40^. The obtained apparent K_d_ values serve as comparative measures between the different experimental conditions. The supershift assay (Supplementary Fig. 2c) was performed under standard EMSA conditions with a final sample volume of 10 µl. The final concentration of *Arc* eRNA 1-200 nt in the eRNA containing samples was 1 µM. The NELF antibody (anti-NELF-E, #ab170104, Abcam) and the DSIF antibody (anti-SPT5, #sc-133217X, Santa Cruz Biotech) were used with the final concentrations of 0.26 µM (2 µl of 1.3 µM was added) and 0.48 µM (2 µl of 2.4 µM), respectively. This corresponds to a ratio of about 2:1 (for NELF) and 4:1 (for DSIF) of the antibody to protein per sample. The NELF titration experiment (Supplementary Fig. 3a) using the four different NELF variants (WT, NELFΔRRM, NELF patch mutant, NELF double mutant) was performed under standard EMSA conditions. NELF variants were added in the following amounts: 0.24, 0.4, 0.8, 1.0, 1.2, 1.6, 2.4 pmol (final concentrations: 0.03, 0.05, 0.1, 0.125, 0.15, 0.2, 0.3 µM).

### Protein-RNA Crosslinking coupled to mass spectrometry

#### Preparation of the paused elongation complex bound to enhancer RNAs

The PEC was prepared as described for EMSAs, except for the fact that the nascent RNA was not radiolabeled and that a 2-fold excess of RNA-DNA hybrid over Pol II was used. 25 µL of 100 µM RNA was annealed with 25 µL of 100 µM template DNA in a total volume of 100 µL (final RNA-DNA hybrid concentration: 25 µM). To assemble the PEC in complex with *Nr4a1*-(b) eRNA (1-100), the following components were added: 1.4 nmol (56 µL of 25 µM) of the RNA-template DNA hybrid, 0.7 nmol (140 µL of 5 µM) of Pol II, 2.8 nmol (28 µL of 100 µM) of non-template DNA, 1.75 nmol (60 µL of 30 µM) DSIF and 1.75 nmol NELF (60 µL of 30 µM). For PEC formation with *Arc* eRNA (1-200), only half of the amount of all these components was used. Following complex formation, the sample was supplemented with 130 µL of 1x complex buffer (20 mM Na-HEPES pH 7.4, 100 mM NaCl, 3 mM MgCl_2_, 1 mM DTT) and additional MgCl_2_ to reach a final MgCl_2_ concentration of 3 mM. Thereafter, 1.75 nmol (67 µL of 26 µM) of *Nr4a1*-(b) eRNA (1-100) was added and incubated for 15 min at 30 °C. To isolate eRNA-bound PEC, the sample was subsequently applied to a Superose 6 10/300 column that had been pre-equilibrated in running buffer (20 mM Na-HEPES pH 7.4, 100 mM NaCl, 3 mM MgCl_2_, 1 mM DTT). The peak representing the eRNA-bound PEC complex was then split into a “main” peak sample and a “shoulder” sample, respectively. Both samples were individually concentrated to 0.5-1 mg/mL using centrifugal filters (100 kDa MWCO, Amicon Ultra-4). They were separately processed during crosslinking mass spec analysis. However, as the detected crosslinking patterns did not differ between the two samples, mass spectrometry data for peak and shoulder samples were merged for data analysis (Supplementary Table 4).

#### Protein-RNA crosslinking and sample preparation for mass spectrometric analysis

Pooled fractions from size exclusion chromatography containing the PEC and either *Arc* eRNA (1-200) or *Nr4a1*-(b) eRNA (1-100) were split in 50 µl aliquots and deposited in 96-well plates. The well plate was put on ice and placed at a distance of approximately 2 cm to the lamps in a Spectrolinker XL-1500 UV Crosslinker (Spectronics Corporation). Samples were irradiated with 3.2 J cm^−2^ of 254 nm UV light in four steps of 0.8 J cm^−2^. Following the crosslinking step, samples were precipitated by adding 3 M sodium acetate (pH 5.2) and ice-cold (−20 °C) ethanol at 0.1 and 3 times the sample volumes, respectively, and incubation at −20 °C overnight. On the following day, the pellet was washed with 80% (v/v) ethanol and dried in vacuum. The samples were dissolved in in 50 µl of 4 M urea in 50 mM Tris (pH 7.9) and further diluted to 1 M urea with 50 mM Tris (pH 7.9). For nuclease digestion, 5 U RNase T1 (ThermoFisher Scientific) and 5 µg of RNase A (Roche Diagnostics) were added per µg of sample, and the solutions were incubated at 52 °C for 2 h. Disulfide bonds in RNase-treated samples were then reduced with tris(2-carboxyethyl)phosphine (2.5 mM final concentration) for 30 min at 37 °C, and free cysteines were alkylated by addition of iodoacetamide (5 mM final concentration) for 30 min at 22 °C in the dark. Following the reduction and alkylation steps, the proteins were digested by addition of sequencing-grade trypsin (Promega) at an enzyme-to-substrate ratio of 1:25. Proteolysis proceeded at 37 °C overnight. Digestion was stopped by addition of 100% formic acid to 2% (v/v) and digested samples were purified by solid-phase extraction using Sep-Pak tC18 cartridges (Waters). The eluate was evaporated to dryness in a vacuum centrifuge and peptide-RNA crosslink products were enriched by titanium dioxide metal oxide affinity chromatography using 10 µm Titansphere PhosTiO beads (GL Sciences). Samples were redissolved in 100 µl of loading buffer (50% acetonitrile, 0.1% trifluoroacetic acid, 10 mg ml^−1^ lactic acid in water) and added to 5 mg of pre-washed TiO_2_ beads and incubated for 30 min on a Thermomixer (Eppendorf) at 1200 rpm to keep the beads in suspension. Beads were settled by centrifugation, the supernatant was removed, and the beads were washed, once with 100 µl of loading buffer and once with 100 µl of washing buffer (loading buffer without lactic acid), by shaking for 15 min, followed by centrifugation steps. Bound peptide-RNA conjugates were eluted from the beads in two steps by adding 75 µl of elution buffer (50 mM diammonium hydrogenphosphate, pH 10.5 adjusted with ammonium hydroxide) using the same procedure (15 min incubation with shaking followed by centrifugation). The eluates were combined and immediately acidified with 100% trifluoroacetic acid to a pH of 2-3. The acidified eluates were purified using Stage tips ^73^ prepared with three plugs of Empore C_18_ disks (3M), and the eluate was again evaporated to dryness. Dried samples were redissolved in water/acetonitrile/formic acid (95:5:0.1, v/v/v) for mass spectrometry analysis.

#### Liquid chromatography-tandem mass spectrometry analysis and identification of protein-RNA crosslinks

LC-MS/MS analysis was performed on an Easy-nLC 1200 HPLC system coupled to an Orbitrap Fusion Lumos mass spectrometer (both ThermoFisher Scientific). Peptides were separated on an Acclaim PepMap RSLC C_18_ column (25 cm × 75 µm, ThemoFisher Scientific) using gradient elution with solvents A (water/acetonitrile/formic acid, 95:5:0.1, v/v/v) and B (water/acetonitrile/formic acid, 20:80:0.1, v/v/v) at a flow rate of 300 nl min^−1^. The gradient was set from 6-40 %B in 60 min. The mass spectrometer was operated in data-dependent acquisition mode using the top speed mode and a cycle time of 3 s. The precursor ion scan was performed in the orbitrap analyzer at a resolution of 120000. MS/MS sequencing was performed on precursors with charge states in the range of +2 to +7, with quadrupole isolation (isolation width 1.2 m/z) and with stepped HCD using normalized collision energies of 23% ±1.15% or 28 ±2.8%. Detection of fragment ions was performed in the orbitrap analyzer at a resolution of 30000 or in the linear ion trap analyzer in rapid mode. Repeated selection of the same precursor m/z was prevented by enabling dynamic exclusion for 30 s after one scan event. For data analysis, the Thermo raw files were converted into mzXML format using msconvert ^74^ and searched using the crosslinking search engine xQuest ^75^ (available from https://gitlab.ethz.ch/leitner_lab/xquest_xprophet). The protein sequence database contained the subunits of the PEC and abundant contaminants identified in the respective samples identified by LC-MS/MS analysis of non-enriched samples. The search only considered the most frequent types of adducts (on all 20 amino acids) with the following nucleotide compositions: A (mass shift = 347.063085 Da), C (323.051851), G (363.057999), U (324.035867), AU (653.088387), CU (629.077153), GU (669.083301), and UU (630.061169), defined as monolinkmw in xquest.def. xQuest search parameters included: enzyme = trypsin, number of missed cleavages = ≤2, MS mass tolerance = ±10 ppm, MS/MS mass tolerance = ±20 ppm (orbitrap detection) or 0.2/0.3 for common and xlink ions (ion trap detection), fixed modification = carbamidomethylation on Cys, variable modification = oxidation of Met. Results were additionally filtered with stricter mass error tolerances according to the experimentally observed mass deviation, a minions value of ≥4, and an xQuest ID score of ≥25. Crosslinking site localizations were directly taken from the xQuest output. All crosslink identifications on PEC subunits are listed in Supplementary Table 4.

### Enhanced UV crosslinking and immunoprecipitation sequencing (eCLIP-seq)

eCLIP-seq was performed as previously described^76^. Briefly, 10 million cells (0, 5, and 25 min KCl treated) were crosslinked (400 mJ/cm^2^ of 254 nM UV light) with a UV crosslinker (Stratagene) that took around 5 min, followed by lysis in lysis buffer (10 mM Tris-HCl [pH 7.6], 300 mM NaCl, 0.1 % Sodium deoxycholate, 1 % Triton X-100, 1 mM EDTA [pH 8.0], 0.5 mM EGTA [pH 8.0], 0.1 % SDS, and protease inhibitors). Samples were carried out with limited digestion with RNase I (AM2294, Ambion) and DNase I (M6101, Promega) to fragment RNAs for 5 min at 25 °C which reactions were stopped by incubated with RNase inhibitor (Y9240, Enzymatics). Two different conditions were lysed in lysis buffers containing straight 0.1 % SDS only or 1 % SDS which was diluted to 0.1 % SDS after 5 min incubation. The lysates were then incubated with antibody (NELF-E; ab170104, Abcam)-conjugated magnetic beads overnight at 4 °C. The immune-complexes were washed with each of the following buffers: high salt buffer (20 mM Tris-HCl [pH 7.6], 1 M NaCl, 1 % Triton X-100, 2 mM EDTA, 0.5% N-lauroylsarcosine, 0.1 % SDS, and 2 M Urea), LiCl buffer (10 mM Tris-HCl [pH 7.6], 250 mM LiCl, 1 % NP-40, 1 % Sodium deoxycholate, 1 mM EDTA, and 2 M Urea), and TET buffer (10 mM Tris-HCl [pH 7.6], 1 mM EDTA, and 0.2 % Tween-20). In each wash, the beads were incubated with wash buffer containing RNase inhibitor for 10 min at 4 °C. RNAs were dephosphorylated with Alkaline Phosphatase (20 U; M0290S, NEB, Ipswich, MA) and T4 PNK (10 U; M0437M, NEB, Ipswich, MA). Subsequently, a 3’ RNA adaptor was ligated onto the RNA with T4 RNA Ligase 1, High Conc. (75 U; M0437M, NEB, Ipswich, MA). Protein-RNA complexes were run on a Bis-Tris SDS-PAGE and the gel containing protein-RNA complexes shifting upwards from the expected size of protein (around 40 up to 150 kD) were cut out. The RNAs were recovered from the gel slices by digesting the protein with proteinase K (8 U; P8107S, NEB, Ipswich, MA) leaving a polypeptide remaining at the crosslinked nucleotide. After precipitation, RNAs were reverse transcribed with SuperScript III Reverse Transcriptase (200 U; 18080-044, ThermoFisher Scientific), the free primer was removed by Exonuclease I (80 U; M0293S, NEB, Ipswich, MA), and a 3’ DNA adaptor was ligated onto the cDNA product with T4 RNA Ligase 1, High Conc. (60 U; M0437M, NEB, Ipswich, MA). Libraries were then amplified with Q5 High-Fidelity Master Mix (M0492S, NEB, Ipswich, MA). Subsequently, PCR products were size-selected in 6 % polyacrylamide TBE gel (210∼350 bp) and purified. The final libraries were sequenced using Illumina NEXTSEQ 500 according to the manufacturer’s instructions. For analysis, adaptor sequences of FASTQ reads were trimmed using cutadapt with ‘-a AGATCGGAAGAGC’ of universal Illumina adapter sequences. The trimmed reads were aligned against a repetitive element-masked mouse genome (mm10) using the STAR aligner. PCR duplicate reads on paired-end (R1 and R2) with same start/end points and same mapped inserts were removed using Sambamba. Then, only R2 reads, which contained information about potential truncation events, were used for further analyses. bamCoverage was used to make visualization tracks. Crosslinking sites were extracted using htseq-clip with ‘-e 2 --site s’ parameter. The very first nucleotides from the eCLIP R2 reads are supposed to represent the crosslinking sites. To examine the distance from the defined 5’ ends of eRNAs and the TSSs to cross-linked sites, we used the ‘bedtools intersect’. Then we calculated the proportion of the crosslinking sites present in six distance windows (up to 200, 201-400, 401-600, 601-800, 801-1,000, or 1,001-2,000 nucleotides) for enhancer and promoter.

### Chromatin Immunoprecipitation (ChIP)

ChIP assays were carried out as previously described with minor modifications^3,17^. At DIV 7, cultured cortical neurons were treated with the indicated conditions, then fixed in crosslinking-buffer (0.1 M NaCl, 1 mM EDTA, 0.5 mM EGTA, 25 mM Hepes-KOH, pH 8.0) containing 1% formaldehyde (252549, Sigma-Aldrich, St. Louis, MO) for 10 min at RT. Crosslinking was quenched by glycine (final 125 mM) for 5 min at RT and harvested in PBS protease inhibitors on ice. Pelleted neurons were lysed in ice-cold buffer I (50 mM HEPES-KOH [pH 7.5], 140 mM NaCl, 1 mM EDTA [pH 8.0], 10 % Glycerol, 0.5 % IGEPAL CA630, and protease inhibitors) to isolate nuclei. Nuclei were sonicated in ice-cold buffer III (10 mM Tris-HCl [pH 8.0], 300 mM NaCl, 0.1 % Sodium deoxycholate, 1 % Triton X-100, 1 mM EDTA [pH 8.0], 0.5 mM EGTA [pH 8.0], and protease inhibitors). The resulting nuclear extracts were centrifuged at 13,200 rpm for 15 min at 4 °C to separate insoluble fraction. The supernatant was then incubated with anti-NELF-A (sc-23599, Santa Cruz Biotech), anti-NELF-E (ab170104, Abcam), or anti-Pol II (N-20) (sc-899X, Santa Cruz Biotech) overnight at 4 °C. Protein A/G PLUS Agarose (sc-2003, Santa Cruz Biotech, Santa Cruz, CA) was added and incubated for 2 hrs at 4 °C. The immune-complexes were pelleted and washed twice with each of the following buffers: low salt buffer (0.1 % SDS, 1 % Triton X-100, 2 mM EDTA, 20 mM Tris-HCl [pH 8.1], 150 mM NaCl), high salt buffer (0.1 % SDS, 1 % Triton X-100, 2 mM EDTA, 20 mM Tris-HCl [pH 8.1], 500 mM NaCl), and LiCl buffer (250 mM LiCl, 1 % IGEPAL CA630, 1 % Sodium deoxycholate, 1 mM EDTA, 10 mM Tris [pH 8.1]). In each wash, the beads were incubated with wash buffer for 10 min at 4 °C. The washed beads were then rinsed once with 1x TE (10 mM Tris-HCl [pH 8.0], 1 mM EDTA). The immune-complexes were eluted from the beads twice by elution buffer (10 mM Tris-HCl [pH 8.0], 1 mM EDTA [pH 8.0], 1 % SDS) at 65 °C for 10 min. The crosslinking was reversed by incubation at 65 °C for 5-6 hrs. The resulting eluate was treated with RNase A (10 μg; Qiagen, Hilden, Germany) for 1 hr at 37 °C and Proteinase K (4 U; P8107S, NEB, Ipswich, MA) for another 2 hr at 55 °C. The DNA was purified by Phenol:Chloroform extraction, followed by PCR purification kit (28106, Qiagen).

### ChIP-seq library construction and data processing

ChIP-seq library construction was performed using NEBNext ChIP-Seq Library Prep Master Mix Set (E6240, NEB, Ipswich, MA) following the manufacturer’s instruction with modifications. Briefly, the end-repaired ChIP DNA fragments were size selected (100∼300 bp), dA-tailed, and then ligated with adaptors. The adaptor-ligated ChIP DNA fragments were digested by USER enzyme and amplified by 14-16 cycles of PCR. The amplified ChIP DNA library was size selected (250∼350 bp) and proceeded to sequencing. ChIP-seq libraries were sequenced on Illumina HISEQ 2500 with 50-bp or NextSeq 500 instrument with 75-bp single-end reads according to manufacturer’s instructions (Illumina) by the UTSW McDermott Next Generation Sequencing Core and Genomics Core. Adapter and low-quality score sequences of FASTQ reads were trimmed using Trim Galore with default parameters. The trimmed FASTQ reads were aligned to UCSC’s mm10 genome using Bowtie2 with default parameters. Reads with mapping quality less than 10 were removed using SAMtools^77^. To normalize the differences in sequencing depths, the mapped reads were “down sampled” to the lowest number of the uniquely mapped reads with duplicates followed by duplicate reads removal using ‘Sambamba’. The bigWig files were generated using ‘bamCoverage’ included in ‘Deeptools’ package for visualization on UCSC genome browser. The coverage values in bigWig files were normalized to RPGC (Reads per genomic content). ChIP peaks were called using MACS with parameters “--gsize mm --broad” against input chromatin samples as control data. To find overlapping peaks under different conditions, we merged the peaks from different samples, called “merged peaks”. Mergepeaks of HOMER was used. To generate ChIP coverage plots, we used HOMER. For HOMER, “makeTagDirectory” program was used. Before making tag directories, we converted BAM files to BED files using BEDTools. We used BED files that contained down-sampled, duplicates removed reads to create tag directories which contain tag information classified per chromosome wise. Using HOMER’s in-built Perl scripts annotatePeaks.pl and analyzeRepeats.pl, data were used to create coverage plots.

### Transcriptional pause release assays on magnetic beads

Pause release assays were adapted from Vos et al., 2018 and performed with a fully complementary scaffold which is similar to the nucleic acid scaffold used for EMSA experiments. The same RNA sequence (25 nt) was used as for EMSA experiments. The template DNA (5’-GAT CAA GCA GTA ATC GTT GCG ATC TGT AGA CTG ACC AAG TTG TCC CGT AGG AGT GTA AGA GAT ATA TGG TAG TAC C; 76 nt) and the non-template DNA (5’-BiotinTEG-GTC TGG TAC TAC CAT ATA TCT CTT ACA CTC CTA CGG GAC AAC TTG GTC AGT CTA CAG ATC GCA ACG ATT ACT GCT TGA TC; 80 nt) sequences were longer than the ones used for the EMSAs in order to allow for the transcription of longer RNA products. Moreover, the non-template DNA had a 4 nt overhang at its 5’-end, which carried a biotin-tag to enable binding to streptavidin magnetic beads. The transcription scaffold contains a 9-base pair (bp) DNA-RNA hybrid, 16 nt of exiting RNA bearing a 5’-^32^P label, 17 nt of upstream DNA and 50 nt of downstream DNA. The radioactively-labeled hybrid of RNA and template DNA was prepared as described for the EMSA experiment. Samples were generally prepared in a *n*-fold master mix (usually a 14x master mix was prepared, which was split into six 2.33x master mixes, from which each master mix was used to produce one time-course transcription experiment). All amounts in the following are related to a 1x mixture. The assembly of Pol II transcription-competent complex originated from the hybrid of RNA and template DNA (2.5 pmol), Pol II (3.75 pmol) and non-template DNA (5 pmol). It was carried out as described for the EMSA experiment, except for the increased amounts of all components. After a final incubation with non-template DNA, the reaction was supplemented with 5x transcription buffer to achieve 1x transcription buffer conditions of 20 mM Na-HEPES pH 7.4, 150 mM NaCl, 3 mM MgCl_2,_ 10 µM ZnCl_2,_ 4% (v/v) glycerol and 2 mM DTT in a final volume of 5 µL. Addition of ATP/CTP/UTP (HTP)-nucleotide mix (1 µL, 100 µM) and incubation at 30°C for 10 min allowed Pol II to transcribe four nucleotides of the implemented G-less cassette before it stalled at +4 position due to GTP omission. Subsequently, DSIF (7.5 pmol) and then NELF (7.5 pmol) were added and each time incubated at 30 °C for 15 min. An *Input* control sample was taken before proceeding to the bead binding step. The master mix (14x) of the assembled PEC complex was then diluted with 1x transcription buffer to a final volume of 180 µL (1.5x of initial bead volume) and was applied to magnetic streptavidin beads (120 µL beads per 14x master mix) (Dynabeads® MyOne™ Streptavidin C1, ThemoFisher Scientific). Prior to this, beads were three-times washed with 1x BW buffer (20 mM Na-HEPES pH 7.5, 100 mM NaCl, 4 % (v/v) glycerol, 0.04% (v/v) Tween20, 0.02% (v/v) IGEPAL CA-630, 2 mM DTT) and finally resuspended in 120 µL 1x transcription buffer. The binding mix (300 µL total volume) was incubated at room temperature on a tube rotator for 20-30 min. After taking off the supernatant, beads with the bound PEC complexes (usually 1/3 of initial RNA was bound, judged by the radioactivity ratio between supernatant and beads) were washed three-times with 1x BW buffer and split into six samples (2.33x of original master mix). The BW buffer was removed, the beads were resuspended in 60 µL of 5 µM eRNA sample in 0.5x RNA buffer (10 mM Na-HEPES pH 7.4, 50 mM KCl, 0.5 mM EDTA) and incubated at room temperature on a tube rotator for 15 min. Subsequently the supernatant was taken off, beads were resuspended in 24 µL of 1x Transcription buffer and the 0 min sample (4.3 µL) was taken and quenched with 2x Stop buffer (6.4 M urea, 50 mM EDTA pH 8.0, 1x TTE buffer). Transcription was allowed to resume by the addition of NTPs (3 µL, 100 µM), the bead mix was immediately returned to a thermomixer and incubated at 30 °C and 900 rpm to avoid sedimentation of the beads. Samples (4.9 µL) were taken after 1, 3, 6, 14 and 25 min and immediately quenched with 2x Stop Buffer (5 µL). Multiple time course experiments were processed in parallel in a phased manner (10 min phasing). Each sample was proteinase K (5 µg) treated at 37 °C for 30 min and then boiled at 90 °C for 4 min to release the bound molecules from the beads. Upon this treatment, samples were separated on a pre-run (30 min at 400 V) denaturing 15 % Urea-PAGE gel (0.5xTTE, 16 x 18 cm x 0.4 mm) for 3 h at 500 V (first 30 min at 400 V). Usually, 4-6 µL of sample were loaded depending on the amount of radioactivity. After the gel run, gels were dried and exposed overnight as described for the EMSA experiments. For a quantification of pause release, the first band above the triple pausing band (as highlighted on the gels in Fig. 4b-d) was densitometrically analyzed. The band intensity was normalized against the total intensity of the corresponding lane and plotted against the time (min).

## Data availability

The Exo-, ChIP- (NELF-A, NELF-E-E and Pan Pol II), eCLIP-, GRO-, mRNA-seq and SHAPE-MaP data from this study are available from NCBI GEO under accession code GSE163113 and GSE139309. ChIP- (H3K27ac, Pol II, and CBP) data was downloaded from GEO (GSE60192 and GSE21161). The mass spectrometry data were deposited to the ProteomeXchange Consortium via the PRIDE partner repository (dataset identifier PXD023298)^78^.

## ACKNOWLEDGEMENTS

We thank Elizabeth Duncan, Olaf Stemmann and Alan Cheung for critically reading the manuscript. We thank Dr. Kunz and Marcel Bowens from the Bayerische Landesanstalt für Landwirtschaft (LfL) for access to pig thymus of unparalleled freshness. We thank Dr. Felix Klatt and Silke Spudeit for technical help, Chris Sarnowski for support with the xQuest software, and we are grateful to the Core Unit Systems Medicine of the University of Würzburg for next-generation sequencing. We thank Patrick Cramer, Seychelle Vos and Carrie Bernecky for providing NELF expression plasmids and for help in establishing the purification of mammalian Pol II. We further thank Birgitta Wöhrl for the DSIF expression plasmid and Norbert Eichner and Gunter Meister for access to a miSeq instrument. This work was supported by the German Research Foundation (DFG, grants KU 3514/1-1 and KU3514/3-1), the Oberfrankenstiftung (P-Nr. 05474), the Elite Network of Bavaria, the University of Bayreuth and the Paul Ehrlich and Ludwig Darmstaedter Prize for Young Researchers (to C.D.K.). This work was also supported by the National Research Foundation of Korea (NRF) grant funded by the Korea government (MSIT), NRF-2019R1A2C2006740, NRF-2019R1A5A6099645, NRF-2017M3A9G7073033 (T.-K.K.), the Brain Research Program of the National Research Foundation (NRF) funded by the Korean government (MSIT), NRF-2020R1I1A1A01067189 (S.-K.K.), NRF-2019M3C7A1031537 (T.-K.K.) and a Simons Foundation Autism Research Initiative-Pilot Award 575147 (T.-K.K.). A.L. acknowledges funding from an ETH Research Grant (ETH-24 16-2) and the ETH Domain Strategic Focus Area “Personalized Health and Related Technologies” (PHRT-503) and would like to thank Paola Picotti (ETH Zurich) for access to instrumentation and infrastructure. The Orbitrap Fusion Lumos mass spectrometer used in this work was purchased using funding from the ETH Scientific Equipment program and the European Union Grant ULTRA-DD (FP7-JTI 115766).

## AUTHOR CONTRIBUTIONS

V.G. and C.-D.K. conceived and designed all Exo-Seq, SHAPE-MaP and *in vitro* experiments. V.G. and F.K. acquired data. A.P. and V.G. performed computational analyses. L.-M.S. helped to establish the pause release assay. T.-K.K. and S.-K.K. designed all experiments related to the characterization of NELF and eRNAs in primary neuron culture. S.-K.K. performed all ChIP-seq, GRO-seq, and eCLIP-seq experiments under the guidance of T.-K.K. D.U. and K.K. performed the bioinformatics analysis of ChIP-seq, GRO-seq, and eCLIP-seq through discussion with T.-K.K. and S.-K.K. M.G. and A.L. performed crosslinking/mass spectrometry experiments and performed data analysis. C.-D.K., T.-K.K. and V.G. drafted the manuscript with input from all authors.

## REFERENCES

1. Deng, W. et al. Controlling Long-Range Genomic Interactions at a Native Locus by Targeted Tethering of a Looping Factor. Cell 149, 1233–1244 (2012).

2. Schier, A. C. & Taatjes, D. J. Structure and mechanism of the RNA polymerase II transcription machinery. Genes Dev 34, 465–488 (2020).

3. Kim, T.-K. et al. Widespread transcription at neuronal activity-regulated enhancers. Nature 465, 182–187 (2010).

4. DeSanta, F. & Natoli, G. A Large Fraction of Extragenic RNA Pol II Transcription Sites Overlap Enhancers. PLoS Biology 8:e1000384 (2010).

5. Li, W. et al. Functional roles of enhancer RNAs for oestrogen-dependent transcriptional activation. Nature 498, 516–520 (2013).

6. Wang, D. et al. Reprogramming transcription by distinct classes of enhancers functionally defined by eRNA. Nature 474, 390–394 (2011).

7. Lai, F. et al. Activating RNAs associate with Mediator to enhance chromatin architecture and transcription. Nature 494, 497–501 (2013).

8. Ørom, U. A. et al. Long Noncoding RNAs with Enhancer-like Function in Human Cells. Cell 143, 46–58 (2010).

9. Mikhaylichenko, O. et al. The degree of enhancer or promoter activity is reflected by the levels and directionality of eRNA transcription. Genes Dev 32, 42–57 (2018).

10. Henriques, T. et al. Widespread transcriptional pausing and elongation control at enhancers. Genes Dev 32, 26–41 (2018).

11. Schwalb, B. et al. TT-seq maps the human transient transcriptome. Science 352, 1225–1228 (2016).

12. Hah, N. et al. A Rapid, Extensive, and Transient Transcriptional Response to Estrogen Signaling in Breast Cancer Cells. Cell 145, 622–634 (2011).

13. Kaikkonen, M. U. et al. Remodeling of the Enhancer Landscape during Macrophage Activation Is Coupled to Enhancer Transcription. Mol Cell 51, 310–325 (2013).

14. Bose, D. A. et al. RNA Binding to CBP Stimulates Histone Acetylation and Transcription. Cell 168, 135–149 (2017).

15. Sigova, A. A. et al. Transcription factor trapping by RNA in gene regulatory elements. Science 350, 978–981 (2015).

16. Rahnamoun, H. et al. RNAs interact with BRD4 to promote enhanced chromatin engagement and transcription activation. Nat Struct Mol Biol 25, 687–697 (2018).

17. Schaukowitch, K. et al. Enhancer RNA Facilitates NELF Release from Immediate Early Genes. Mol Cell 56, 29–42 (2014).

18. Danko, C. G. et al. Signaling Pathways Differentially Affect RNA Polymerase II Initiation, Pausing, and Elongation Rate in Cells. Mol Cell 50, 212–222 (2013).

19. Malik, A. N. et al. Genome-wide identification and characterization of functional neuronal activity–dependent enhancers. Nature 17, 1330–1339 (2014).

20. Afik, S. et al. Defining the 5΄ and 3΄ landscape of the Drosophila transcriptome with Exo-seq and RNaseH-seq. Nucleic Acids Res 45, e95 (2017).

21. Qian, X., Zhao, J., Yeung, P. Y., Zhang, Q. C. & Kwok, C. K. Revealing lncRNA Structures and Interactions by Sequencing-Based Approaches. Trends Biochem Sci 44, 33–52 (2019).

22. Andersson, R. et al. An atlas of active enhancers across human cell types and tissues. Nature 507, 455–461 (2014).

23. Lucks, J. B. et al. Multiplexed RNA structure characterization with selective 2ʹ-hydroxyl acylation analyzed by primer extension sequencing (SHAPE-Seq). Proc Natl Acad Sci USA 108, 11063–11068 (2011).

24. Siegfried, N. A., Busan, S., Rice, G. M., Nelson, J. A. E. & Weeks, K. M. RNA motif discovery by SHAPE and mutational profiling (SHAPE-MaP). Nat Methods 11, 959–965 (2014).

25. Smola, M. J., Rice, G. M., Busan, S., Siegfried, N. A. & Weeks, K. M. Selective 2ʹ-hydroxyl acylation analyzed by primer extension and mutational profiling (SHAPE-MaP) for direct, versatile and accurate RNA structure analysis. Nat Protoc 10, 1643–1669 (2015).

26. Busan, S. & Weeks, K. M. Accurate detection of chemical modifications in RNA by mutational profiling (MaP) with ShapeMapper 2. RNA 24, 143–148 (2018).

27. Reuter, J. S. & Mathews, D. H. RNAstructure: software for RNA secondary structure prediction and analysis. BMC Bioinformatics 11, 129 (2010).

28. Vos, S. M., Farnung, L., Urlaub, H. & Cramer, P. Structure of paused transcription complex Pol II-DSIF-NELF. Nature 560, 601–606 (2018).

29. Bernecky, C., Herzog, F., Baumeister, W., Plitzko, J. M. & Cramer, P. Structure of transcribing mammalian RNA polymerase II. Nature 529, 551–554 (2016).

30. Missra, A. & Gilmour, D. S. Interactions between DSIF (DRB sensitivity inducing factor), NELF (negative elongation factor), and the Drosophila RNA polymerase II transcription elongation complex. Proc Natl Acad Sci USA 107, 11301–6 (2010).

31. Fujinaga, K. et al. Dynamics of Human Immunodeficiency Virus Transcription: P-TEFb Phosphorylates RD and Dissociates Negative Effectors from the Transactivation Response Element. Mol Cell Biol 24, 787–795 (2003).

32. Peterlin, B. M. & Price, D. H. Controlling the Elongation Phase of Transcription with P-TEFb. Mol Cell 23, 297–305 (2006).

33. Yamada, T. et al. P-TEFb-Mediated Phosphorylation of hSpt5 C-Terminal Repeats Is Critical for Processive Transcription Elongation. Mol Cell 21, 227–237 (2006).

34. Leppek, K., Das, R. & Barna, M. Functional 5ʹ UTR mRNA structures in eukaryotic translation regulation and how to find them. Nat Rev Mol Cell Biol 19, 158–174 (2018).

35. Yamaguchi, Y. et al. NELF, a Multisubunit Complex Containing RD, Cooperates with DSIF to Repress RNA Polymerase II Elongation. Cell 97, 41–51 (1999).

36. Rao, J. N. et al. Structural studies on the RNA-recognition motif of NELF E, a cellular negative transcription elongation factor involved in the regulation of HIV transcription. Biochemical J 400, 449–456 (2006).

37. Rao, J. N., Schweimer, K., Wenzel, S., Wöhrl, B. M. & Rösch, P. NELF-E RRM Undergoes Major Structural Changes in Flexible Protein Regions on Target RNA Binding. Biochemistry 47, 3756–3761 (2008).

38. Yamaguchi, Y., Inukai, N., Narita, T., Wada, T. & Handa, H. Evidence that Negative Elongation Factor Represses Transcription Elongation through Binding to a DRB Sensitivity-Inducing Factor/RNA Polymerase II Complex and RNA. Mol Cell Biol 22, 2918–2927 (2002).

39. Pagano, J. M. et al. Defining NELF-E RNA Binding in HIV-1 and Promoter-Proximal Pause Regions. PLoS Genetics 10, e1004090–11 (2014).

40. Vos, S. M. et al. Architecture and RNA binding of the human negative elongation factor. eLife 5, e14981–27 (2016).

41. Zuber, P. K. et al. Structure and nucleic acid binding properties of KOW domains 4 and 6-7 of human transcription elongation factor DSIF. Sci Rep 8, 11660 (2018).

42. Lyskov, S. et al. Serverification of Molecular Modeling Applications: The Rosetta Online Server That Includes Everyone (ROSIE). Plos One 8, e63906 (2013).

43. Watkins, A. M., Rangan, R. & Das, R. FARFAR2: Improved De Novo Rosetta Prediction of Complex Global RNA Folds. Structure 28, 963–976.e6 (2020).

44. Cheng, B. & Price, D. H. Analysis of factor interactions with RNA polymerase II elongation complexes using a new electrophoretic mobility shift assay. Nucleic Acids Res 36, e135–e135 (2008).

45. Mohammed, H. et al. Endogenous Purification Reveals GREB1 as a Key Estrogen Receptor Regulatory Factor. Cell Rep 3, 342–349 (2013).

46. Dengl, S. & Cramer, P. Torpedo Nuclease Rat1 Is Insufficient to Terminate RNA Polymerase II in Vitro. J Biol Chem 284, 21270–21279 (2009).

47. Cheng, B. & Price, D. H. Properties of RNA Polymerase II Elongation Complexes Before and After the P-TEFb-mediated Transition into Productive Elongation. J Biol Chem 282, 21901–21912 (2007).

48. Arner, E. et al. Transcribed enhancers lead waves of coordinated transcription in transitioning mammalian cells. Science 347, 1010–1014 (2015).

49. Adelman, K. & Lis, J. T. Promoter-proximal pausing of RNA polymerase II: emerging roles in metazoans. Nat Rev Genet 13, 720–731 (2012).

50. Davidovich, C., Zheng, L., Goodrich, K. J. & Cech, T. R. Promiscuous RNA binding by Polycomb repressive complex 2. Nat Struct Mol Biol 20, 1250–1257 (2013).

51. Beltran, M. et al. The interaction of PRC2 with RNA or chromatin is mutually antagonistic. Genome Res 26, 896–907 (2016).

52. Wang, X. et al. Molecular analysis of PRC2 recruitment to DNA in chromatin and its inhibition by RNA. Nat Struct Mol Biol 24, 1028–1038 (2017).

53. Beltran, M. et al. G-tract RNA removes Polycomb repressive complex 2 from genes. Nat Struct Mol Biol 26, 899–909 (2019).

54. Wang, X. et al. Targeting of Polycomb Repressive Complex 2 to RNA by Short Repeats of Consecutive Guanines. Mol Cell 65, 1056–1067.e5 (2017).

55. Cramer, P. Organization and regulation of gene transcription. Nature 573, 45–54 (2019).

56. Hnisz, D., Shrinivas, K., Young, R. A., Chakraborty, A. K. & Sharp, P. A. A Phase Separation Model for Transcriptional Control. Cell 169, 13–23 (2017).

57. Molliex, A. et al. Phase Separation by Low Complexity Domains Promotes Stress Granule Assembly and Drives Pathological Fibrillization. Cell 163, 123–133 (2015).

58. Castello, A. et al. Comprehensive Identification of RNA-Binding Domains in Human Cells. Mol Cell 63, 696–710 (2016).

59. Henninger, J. E. et al. RNA-Mediated Feedback Control of Transcriptional Condensates. Cell 184, 207–225 (2021).

60. Shii, L., Song, L., Maurer, K., Zhang, Z. & Sullivan, K. E. SERPINB2 is regulated by dynamic interactions with pause-release proteins and enhancer RNAs. Mol Immunol 88, 20–31 (2017).

61. Hsieh, C.-L. et al. Enhancer RNAs participate in androgen receptor-driven looping that selectively enhances gene activation. Proc National Acad Sci USA 111, 7319–7324 (2014).

62. Kim, T.-K. & Shiekhattar, R. Architectural and Functional Commonalities between Enhancers and Promoters. Cell 162, 948–959 (2015).

63. Hah, N., Murakami, S., Nagari, A., Danko, C. G. & Kraus, W. L. Enhancer transcripts mark active estrogen receptor binding sites. Genome Res 23, 1210–1223 (2013).

64. Lam, M. T. Y. et al. Rev-Erbs repress macrophage gene expression by inhibiting enhancer-directed transcription. Nature 498, 511–515 (2013).

65. Core, L. J., Waterfall, J. J. & Lis, J. T. Nascent RNA Sequencing Reveals Widespread Pausing and Divergent Initiation at Human Promoters. Science 322, 1845–1848 (2008).

66. Cazenave, C. & Uhlenbeck, O. C. RNA template-directed RNA synthesis by T7 RNA polymerase. Proc Natl Acad Sci USA 91, 6972–6976 (1994).

67. Brunelle, J. L. & Green, R. Chapter Five In Vitro Transcription from Plasmid or PCR-amplified DNA. Methods Enzymol 530, 101–114 (2013).

68. Petrov, A., Wu, T., Puglisi, E. V. & Puglisi, J. D. Chapter Seventeen RNA Purification by Preparative Polyacrylamide Gel Electrophoresis. Methods Enzymol 530, 315–330 (2013).

69. Merino, E. J., Wilkinson, K. A., Coughlan, J. L. & Weeks, K. M. RNA Structure Analysis at Single Nucleotide Resolution by Selective 2‘-Hydroxyl Acylation and Primer Extension (SHAPE). J Am Chem Soc 127, 4223–4231 (2005).

70. Liu, F., Somarowthu, S. & Pyle, A. M. Visualizing the secondary and tertiary architectural domains of lncRNA RepA. Nat Chem Biol 13, 282–289 (2017).

71. Bernecky, C., Plitzko, J. M. & Cramer, P. Structure of a transcribing RNA polymerase II-DSIF complex reveals a multidentate DNA-RNA clamp. Nat Struct Mol Biol 24, 809–815 (2017).

72. Berger, I., Fitzgerald, D. J. & Richmond, T. J. Baculovirus expression system for heterologous multiprotein complexes. Nat Biotechnol 22, 1583–1587 (2004).

73. Rappsilber, J., Ishihama, Y. & Mann, M. Stop and Go Extraction Tips for Matrix-Assisted Laser Desorption/Ionization, Nanoelectrospray, and LC/MS Sample Pretreatment in Proteomics. Anal Chem 75, 663–670 (2003).

74. Chambers, M. C. et al. A cross-platform toolkit for mass spectrometry and proteomics. Nat Biotechnol 30, 918–920 (2012).

75. Walzthoeni, T. et al. False discovery rate estimation for cross-linked peptides identified by mass spectrometry. Nat Methods 9, 901–903 (2012).

76. Nostrand, E. L. V. et al. Robust transcriptome-wide discovery of RNA-binding protein binding sites with enhanced CLIP (eCLIP). Nat Methods 13, 508–514 (2016).

77. Li, H. et al. The Sequence Alignment/Map format and SAMtools. Bioinformatics 25, 2078–2079 (2009).

78. Perez-Riverol, Y. et al. The PRIDE database and related tools and resources in 2019: improving support for quantification data. Nucleic Acids Res 47, D442–D450 (2019).

