## Supplementary_Information for "Enhancer RNAs stimulate Pol II pause release by harnessing multivalent interactions to NELF"

**Supplementary Information includes:**

8 Supplementary Figures

4 Supplementary Tables

### Supplementary Fig. 1

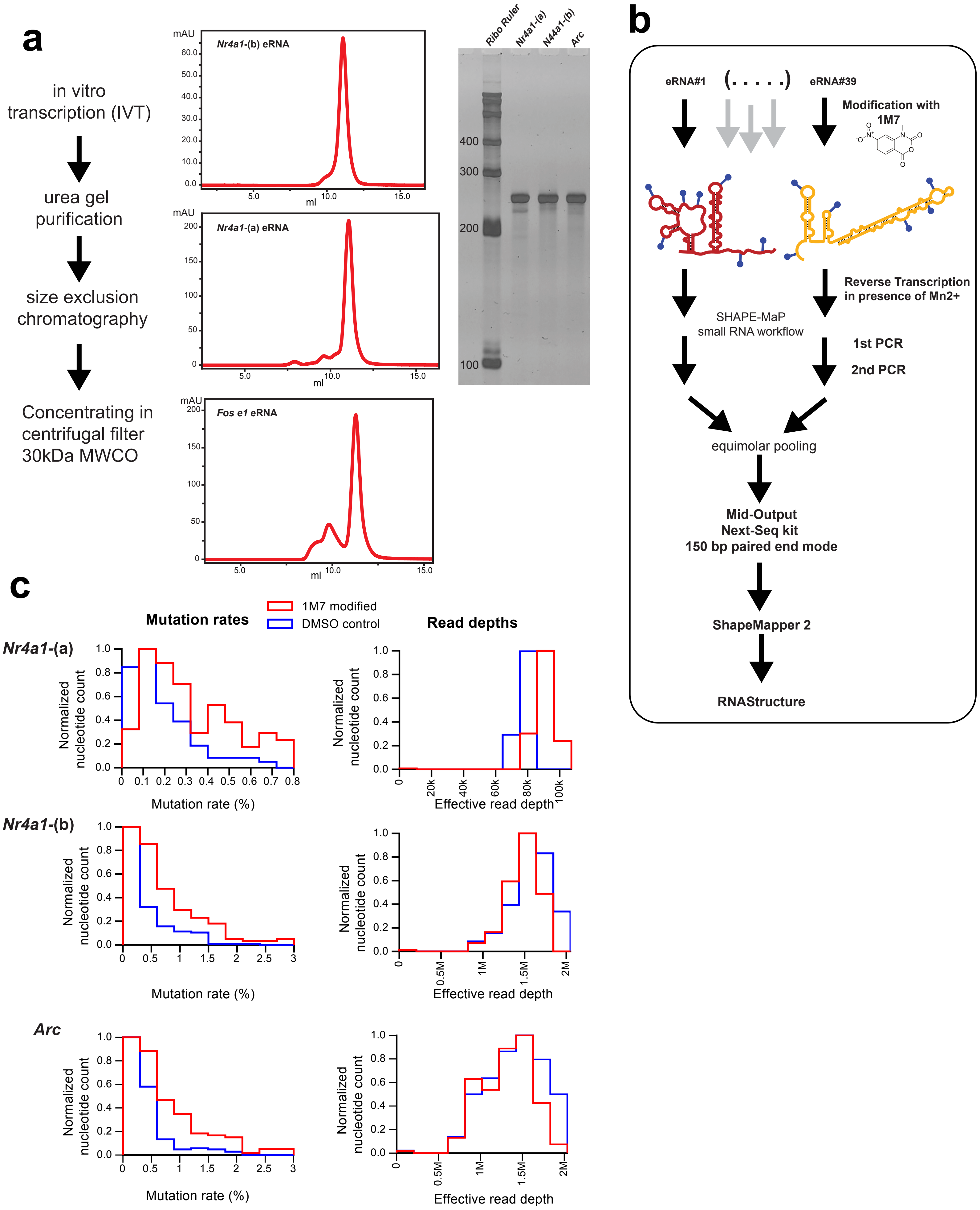

**Supplementary Fig. 1 | SHAPE-MaP workflow and mapping statistics. a,** Workflow of eRNA production for SHAPE-MaP. Representative gel filtration chromatograms of eRNAs and an analytical urea PAGE of purified *Nr4a1*-(a), *Nr4a1*-(b) and *Arc* eRNAs. **b,** Workflow of SHAPE-MAP library construction and analysis. **c,** Mutation rates and read depth statistics for *Arc*, *Nr4a1*-(a) and *Nr4a1*-(b) eRNAs. The elevated mutation rate for the 1M7-treated eRNAs (red line) above the background mutation rate for the DMSO control (blue line) and the corresponding read depth profile and including the effective read depth (shown in lighter colors) confirm the high quality of our SHAPE-MaP data.

**Supplementary Fig. 2**

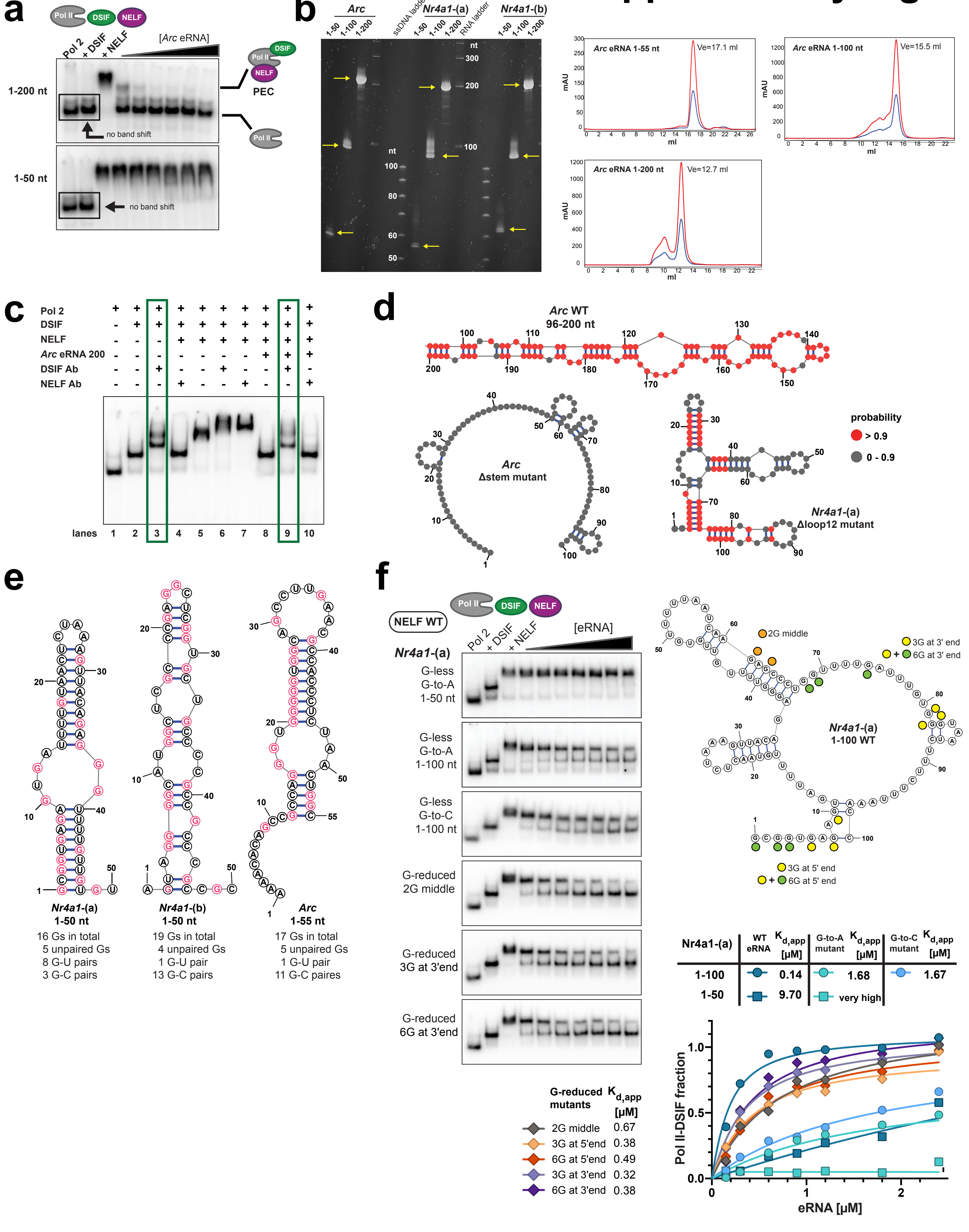

**Supplementary Fig. 2 | EMSA experiments demonstrate that eRNAs trigger NELF release from the paused elongation complex (PEC). a,** EMSA performed on a transcription bubble with a shorter, 15 nt long, nascent RNA (*versus* 25 nt long in Fig. 2b-d). Addition of DSIF does not lead to a shift of the mobility of the initial band that contains Pol II bound to the transcription bubble. Only utilization of a longer nascent RNA (25 nt; see Fig. 2b-d) leads to a visible shift upon DSIF addition, which confirms the binding of DSIF to Pol II. **b,** Quality assessment of *in vitro* produced eRNAs that were used for functional assays. An analytical urea PAGE for *Arc*, *Nr4a1-(a)* and *-(b)* fragments (1-50, 1-100 and 1-200) is shown. The right panel shows representative size exclusion chromatography elution profiles of exemplary *Arc* eRNA fragments. **c,** An EMSA supershift assay confirms the identity of the observed EMSA bands. To that end anti-Strep antibodies and NELF-E antibodies were used (Spt5, the large subunit of DSIF, carries a Strep-tag in our experimental setup). Only the Strep-tag antibody against DSIF (lane 9) and not the NELF-E antibody (lane 10) is able to shift the band after *Arc* 1-200 nt eRNA addition (lane 8). **d,** Secondary structures of the wild type *Arc* eRNA (96-200) fragment (length = 104 nt), the *Arc* eRNA $\Delta$stem mutant (length = 100 nt) and the *Nr4a1*-(a) $\Delta$loop 12 mutant (length = 102 nt) are shown. See also Supplementary Fig. 5b. EMSA conditions were analogous to Fig. 2b. **e,** Secondary structures of *Nr4a1*-(a), -(b) 1-50 and *Arc* 1-55 nt fragment predicted with *RNAStructure*. The structure of *Arc* eRNA (1-55) is consistent with the structure within the 1-200 nt fragment determined by SHAPE-MaP. All guanosines are highlighted in pink. The total G content between the fragments is similar (16-19 Gs). While most of the Gs in *Nr4a1*-(b) and *Arc* are paired with Cs, Gs in *Nr4a1*-(a) are located in loop regions or G-U pairs. **f,** The left panel shows EMSAs performed with G-less *Nr4a1*-(a) 1-50, 1-100 fragments. In these, all guanosines (16 guanosines for the 1-50 fragment and 25 guanosines for the 1-100 fragment) were substituted with adenosine (G-to-A mutant) or cytidine (G-to-C mutant), or different G-reduced *Nr4a1*-(a) (1-100) mutants, where two guanosines (2G middle; G61/63), three guanosines (3G at 3'end: G68/69/74) or six guanosines (6G at 3'end; G68/69/74/81/82/83) had been restored. In the right panel the restored guanosines are highlighted along the secondary structure of the wild type *Nr4a1*-(a) 1-100 fragment, which is based on our SHAPE-MaP structure for the 1-200 fragment. Below, the quantification of NELF release for all G-mutants shown on the left plus two additional mutants (EMSA gels not shown; 3G at 5'end: G8/10/12; 6G at 5'end: G3/4/6/8/10/12) is shown (see Fig. 2e). The quantification of wild-type fragments is added to aid comparison (see Fig. 2b).

### Supplementary Fig. 3

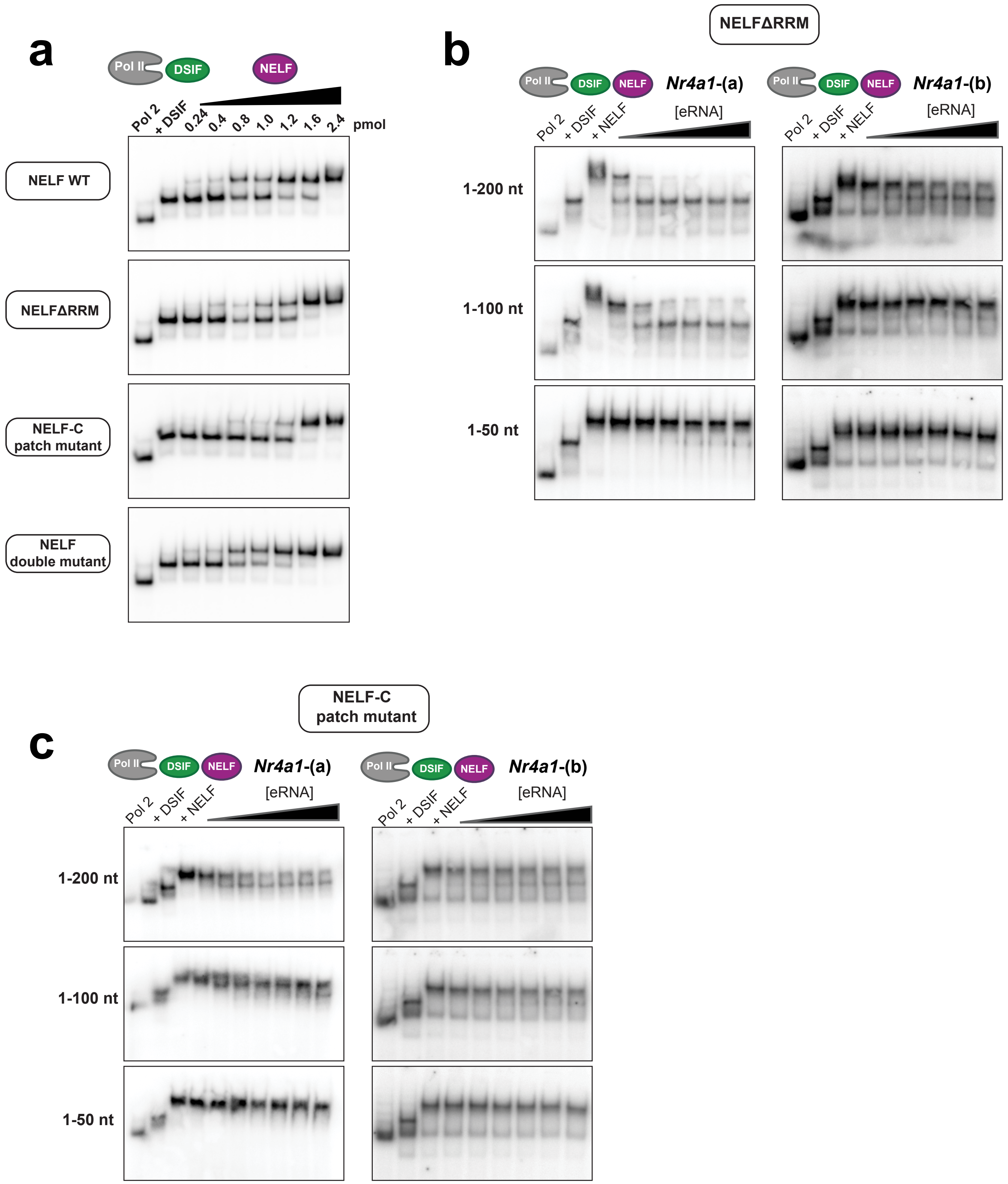

**Supplementary Fig. 3 | NELF mutants form the PEC as efficiently as wild-type NELF, however, their eRNA-driven detachment from Pol II is dramatically reduced. a,** An EMSA titration experiment of WT NELF and different NELF mutants (NELFΔRRM, patch mutant and the double mutant) to preformed Pol II-DSIF complexes demonstrates that wild type and mutant NELF variants form the PEC comparably well. EMSAs performed with the fragments 1-50, 1-100 and 1-200 of *Nr4a1*-(a) and *Nr4a1*-(b) in the presence of either the NELFΔRRM mutant **(b)**, or the NELF patch mutant **(c)**. EMSAs were performed as described in Fig. 2b.

### Supplementary Fig. 4

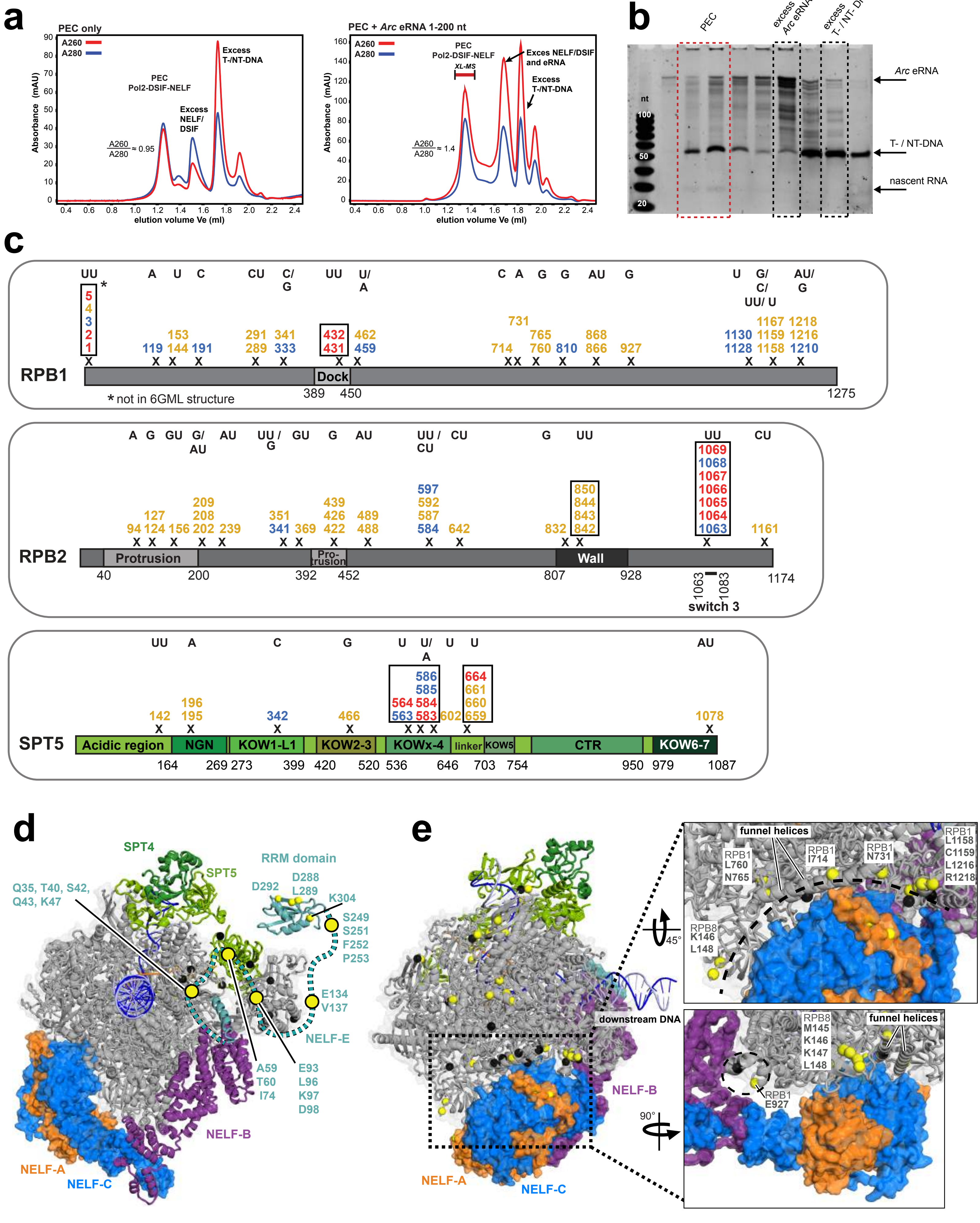

**Supplementary Fig. 4 | UV-crosslinking mass spectrometry of an eRNA-bound PEC suggests that eRNAs detach NELF from Pol II by interfering with the NELF-Pol II binding interface. a,** Elution profiles of the PEC assembly without added eRNA (chromatogram on the left) and with added *Arc* eRNA (1-200) (chromatogram on the right) on an analytical Superose 6 3.2/300 Increase column. Fractions from a preparative Superose 6 column corresponding to PEC peak (marked XL-MS) were used for UV crosslinking coupled to MS/MS experiments to determine eRNA binding sites on the PEC. An increased ratio of A260/A280 absorbance observed in the PEC+eRNA chromatogram compared to the PEC only control indicates eRNA binding to the PEC. **b,** Analytical urea gel of size exclusion chromatography fractions of the experiment in (a) using *Arc* eRNA (1-200). Fractions containing the PEC show the presence of T-DNA, NT-DNA, nascent RNA and *Arc* eRNA. **c,** Positions of RNA-crosslinked residues (Score > 28) on the mammalian Pol II subunits RPB1 and RPB2, as well on the DSIF subunit SPT5. Crosslinks for the experiment with *Arc* eRNA 1-200 fragment are shown in blue, crosslinks for *Nr4a1*-(b) eRNA (1-100) are in orange and residues found in both experiments are highlighted in red. To aid visualization crosslinked residues with a distance of +/- 15 residues were clustered together. Each protein is depicted as a linear bar, relevant domains are indicated. The crosslinked RNA fragments are indicated above the crosslinked amino acids. Nascent RNA crosslinks are boxed. **d,** RNA crosslinks to the NELF-E RRM domain (PDB code 2JX2) and to the structurally unresolved parts of NELF- E (shown as a dashed line) are shown in context of the entire PEC (PDB code 6GML). Crosslinked residues in NELF-E are highlighted in yellow. Residues on Pol II and DSIF previously shown to form lysine-lysine crosslinks with NELF-E along its unresolved parts are shown as black spheres. **e,** RNA-crosslinked residues on Pol II (shown as yellow spheres) are located in the interface between Pol II and the NELF-AC lobe. Previously reported residues on Pol II that form lysine-lysine crosslinks to the NELF-AC lobe are shown as black spheres (PDB: 6GML).

### Supplementary Fig. 5

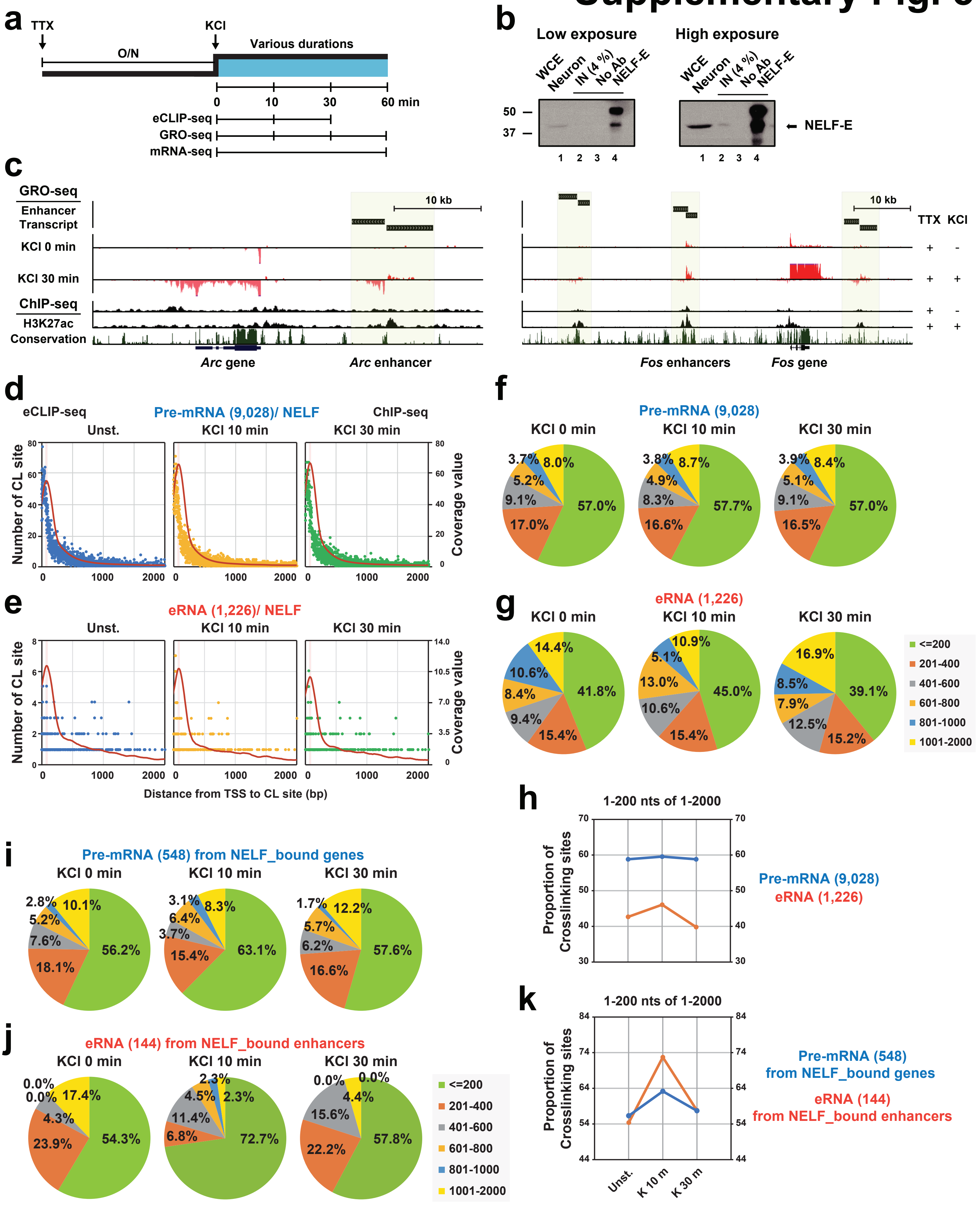

**Supplementary Fig. 5 | eCLIP reveals that NELF interacts with the 5'-end regions of neuronal pre-mRNAs and eRNAs. a,** Experimental scheme for eCLIP-seq, GRO-seq, and mRNA-seq *in vivo*. **b,** Western in two different exposures showing the amount of NELF-E pulled down under eCLIP condition. **c,** Example tracks of *de novo* transcript calling on enhancers from GRO-seq that was identified based on the H3K27ac enriched peaks at the activity-induced genes, *Arc* and *Fos* gene loci. (**d** and **e**) The distribution of the crosslinking sites and NELF ChIP-seq coverage profiles for for total annotated pre-mRNAs (9,028) (**d**) and total intergenic eRNAs (1,226) (**e**) at different time points after KCl stimulation – same as Fig. 5b,c except showing a larger downstream region (up to 2 kb). Each dot indicates the location of the crosslinking site. Blue, yellow, and green indicates 0, 10, and 30 min KCl, respectively. (**f**, **g**, **i** and **j**) Pie chart showing the proportion of the crosslinking sites present in six distance windows (1-200, 201-400, 401-600, 601-800, 801-1,000 or 1,001-2,000 nts) for total annotated (9,028) (**f**) or KCl-up/ NELF_bound (548) (**I**) pre-mRNAs and total intergenic (1,226) (**g**) or KCl-up/ NELF_bound (144) (**J**) eRNAs at different time points after KCl stimulation. (**h** and **k**) Proportion of the crosslinking sites for 1-200 nts of 1-2,000 downstream region for total pre-mRNA (9,028) or eRNA (1,226) (**h**) and NELF bound pre-mRNA (548) or eRNA (144) (**k**).

### Supplementary Fig. 6

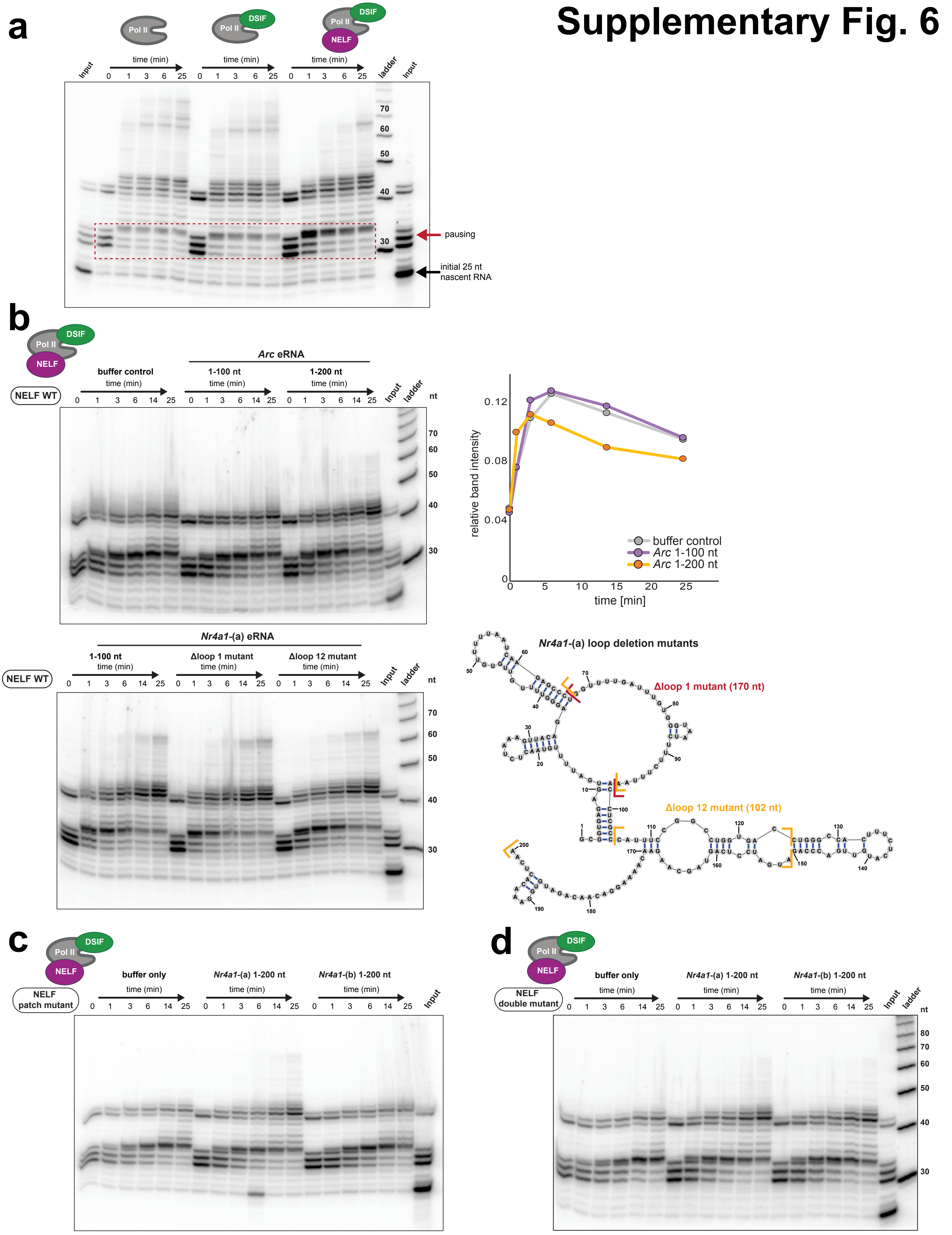

**Supplementary Fig. 6 | DSIF and NELF-dependent pause stabilization and supplementary pause release assays.** All assays were performed as described for Fig. 6a,b. **a,** Transcription assay that verifies pause stabilization in presence of both DSIF and NELF. **b,** Pause release assay with WT NELF and *Arc* (1-100) and (1-200) (top gel) or *Nr4a1*-(a) mutants (bottom gel) (related to Fig. 6b). Next to the top gel the corresponding quantification of the pause release is shown. The quantification for the bottom gel part of Fig. 6b. The secondary structure of *Nr4a1*-(a) (1-200) is shown to the right of the top gel and the deleted regions in the *Nr4a1*-(a) $\Delta$loop 1 (red) or the $\Delta$loop 12 mutant (orange) are marked in the structure. **c,** Pause release assay with the NELF patch mutant (related to Fig. 6c). **d,** Pause release assay using the NELF double mutant (related to Fig. 6d).

### Supplementary Fig. 7

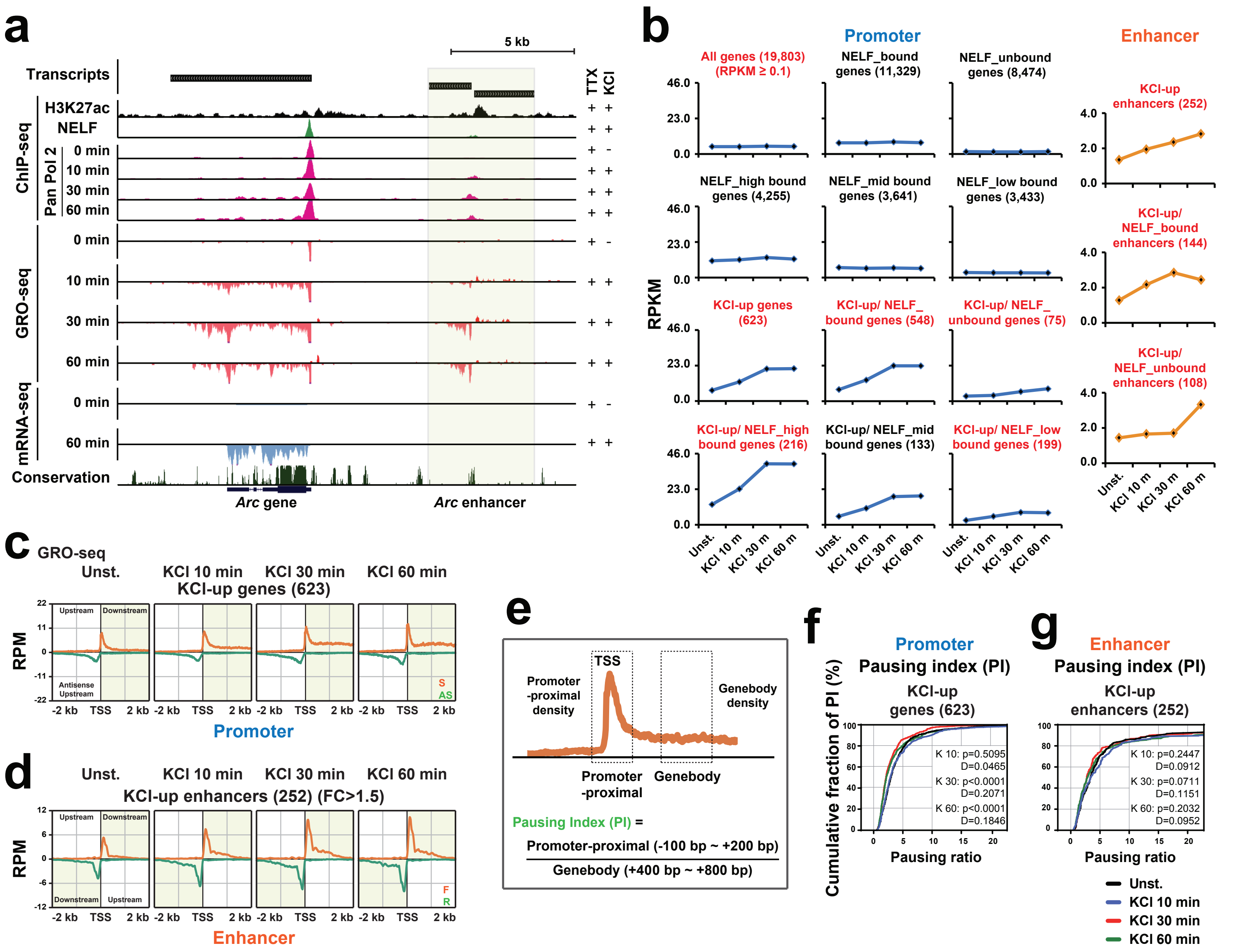

**Supplementary Fig. 7 | NELF-dependent promoter-proximal pausing at neuronal activity-induced genes strongly correlates with the induction levels of these genes. a,** Further example of activity-induced gene, *Arc* and nearby enhancer (shaded area) (related to Fig. 7a). **b**, Transcription induction profiles of pre-mRNAs and eRNAs determined based on GRO-seq data seq in response to 0, 10, 30 and 60 min KCl treatment. All different categories of genes are shown (related to Fig. 7b). (**c** and **d**) The average GRO-seq profiles at 623 KCl-up genes (**c**) and 252 KCl-up enhancers (FC>1.5) (**d**) (related to Fig. 7c,d). The profiles are described at ±2 kb region centered on the TSSs or the enhancer centers upon 0, 10, 30 and 60 min KCl stimulations. Orange line denotes sense strand at promoters and forward strand at enhancers. Green line denotes anti-sense at promoters and reverse at enhancers. **e,** An illustration for calculation of Pausing index (PI) by GRO-seq transcripts at promoter-proximal vs genebody. The promoter-proximal is defined from 100 bp upstream to 200 bp downstream of the TSS. The genebody is defined from 400 to 800 bp downstream of the TSS. (**f** and **g**) PI profiles of 623 KCl-up genes (**f**) and 252 KCl-up enhancers (**g**). This gene group is analyzed for 4 time points (Un, 10, 30 and 60 min KCl). Statistical significance between cumulative probability graphs was determined by the Kolmogorov-Smirnov test.

### Supplementary Fig. 8

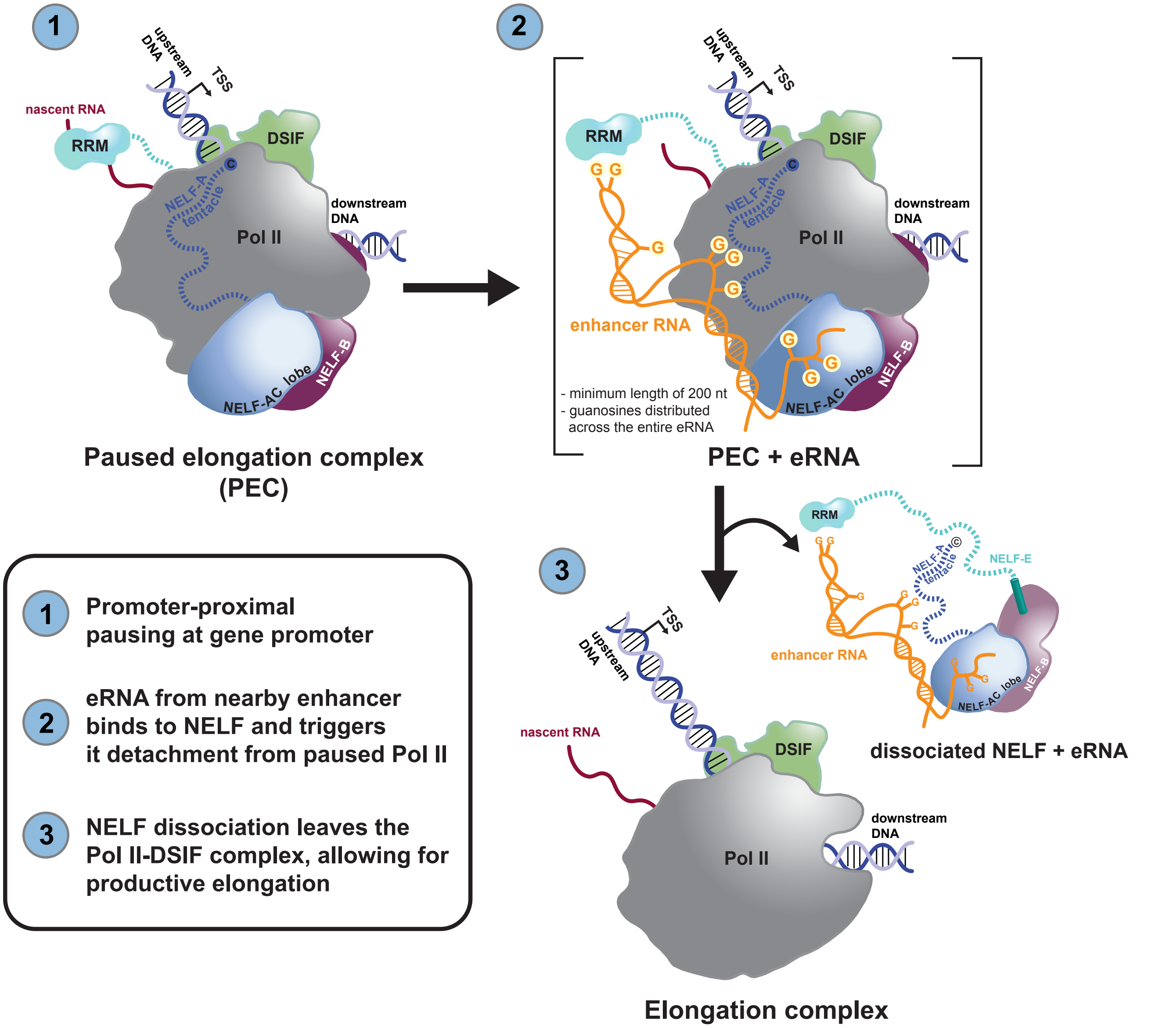

**Supplementary Fig. 8 | Model of how multivalent, allosteric interactions between an eRNA and the paused elongation complex (PEC) stimulate Pol II pause release.** Following the transcription of a nearby, activated enhancer, an eRNA molecule contacts the paused elongation complex (consisting of Pol II, DSIF and NELF) at the target gene promoter. The eRNA interacts with NELF at multiple sites (positive patches on NELF-AC, the NELF-A tentacle and the NELF-E RRM domain). These multivalent, allosteric interactions collectively trigger NELF detachment from the PEC. The resulting elongation complex (consisting of Pol II and DSIF) then resumes transcription elongation. Unpaired guanosines within an eRNA are essential for its ability to detach NELF from the PEC.
